# Antibody-Antigen Affinity Prediction with Chain-Aware Protein Language Modeling

**DOI:** 10.64898/2026.06.19.733375

**Authors:** Harshit Singh, Aastha Malhotra, Satya Pratik Srivastava, Rajeev Kumar Singh, Rohan Gorantla

## Abstract

**Motivation:** Antibody-antigen affinity determines which antibodies advance in therapeutic discovery, repertoire analysis and affinity maturation, but experimental measurements are sparse relative to the scale of sequence libraries. Structure-based predictors can exploit interface geometry when reliable complexes are available, yet early discovery often requires ranking many heavy-light chain pairs against antigens for which no complex structure exists. Existing sequence-based models are scalable, but frequently compress heavy and light chains into a single antibody representation or concatenate antibody and antigen features, potentially obscuring the chain-specific and epitope-specific signals that drive binding.

**Results:** We present AbAffinity, a sequence-only chain-aware three-stream architecture that maintains heavy chain, light chain and antigen as distinct streams. It integrates frozen ESM-2 embeddings with heavy-chain CDR-focused pooling, heavy-light self-attention, adaptive fusion and gated cross-attention, training only a compact interaction module. Under nested ten-fold cross-validation on SAAINT-DB, AbAffinity achieves Pearson *r* = 0.838 ± 0.033, Spearman *ρ* = 0.828 ± 0.027 and RMSE = 0.738 ± 0.078 pK_d_. AbAffinity maintains meaningful performance on stricter PDB-disjoint and antigen sequence-identity splits, and held-out complexes. Evaluation on external natural-complex benchmarks further supports consistent transfer. Partner-shuffling, attribution and uncertainty analyses support partner-conditioned prediction and biologically plausible residue-level hypotheses. AbAffinity provides a lightweight, explainable sequence-first framework for antibody triage when structural information is limited or unavailable.

## 1 Introduction

Antibody-antigen binding affinity is a central determinant of therapeutic antibody discovery, vaccine immunology, B-cell receptor (BCR) repertoire analysis and affinity maturation. In each of these settings, the experimental bottleneck is scale: large numbers of antibody sequences can be generated, observed or designed, but only a small fraction can be synthesised, structurally characterised or assayed. Computational affinity prediction can therefore reduce screening burden by prioritising promising binders, guiding mutational design and helping interpret immune repertoires before costly wet-lab validation (Lu et al., 2020; Zhang et al., 2022; Zhao et al., 2025). For sequence-based affinity prediction to be useful in early discovery, a model must generalise beyond closely related antibody-antigen interactions, scale efficiently to large antibody, BCR and mutational libraries, and provide residue-level rationales that support experimental follow-up.

Structure-based approaches provide an important route to affinity prediction when high-quality antibody-antigen complexes are available. Docking poses, interface geometry, solvent exposure and contact descriptors can capture physical determinants of binding and have supported systematic studies of antibody recognition (Dunbar et al., 2014; Guest et al., 2021b; Myung et al., 2022; Liu et al., 2014). Curated resources such as AB-Bind, SKEMPI 2.0 and more recent structural databases have expanded benchmarking of binding and mutation effects, while also highlighting the limited size, heterogeneity and assay dependence of experimentally measured affinity data (Sirin et al., 2016; Jankauskaité et al., 2019; Huang et al., 2025). However, structure-based workflows remain difficult to deploy at the scale required in early discovery, where millions of sequences may need to be ranked and reliable complex structures are often unavailable. Moreover, structural availability does not by itself guarantee accurate affinity calibration, particularly when coarse interface descriptors fail to capture sequence context, partner specificity or assay-dependent energetic effects. These limitations motivate complementary sequence-first methods that can be applied before structural refinement.

Sequence-based models are attractive because they scale naturally to large libraries and can exploit protein language models that encode evolutionary, structural and biochemical regularities from large protein corpora (Elnaggar et al., 2021; Ruffolo et al., 2021; Lin et al., 2023; Burbach and Briney, 2024; Unsal et al., 2022). Recent affinity predictors, including DG-Affinity, MVSF-AB, MINT and related antibody, protein-protein and drug-target models, show that useful binding signal can be extracted from sequence embeddings (Öztürk et al., 2018; Yugandhar and Gromiha, 2014; Abbasi et al., 2020; Chen et al., 2019; Jin et al., 2024; Wang et al., 2024; Ullanat et al., 2026; Gurusinghe et al., 2025; Gorantla et al., 2025; Gorantla, 2025). Yet several architectural limitations remain. Many models combine antibody and antigen features through concatenation, shallow attention or global pooling, which can obscure the partner-specific contributions that determine binding (Xu et al., 2022; Gorantla et al., 2024b). Others compress the antibody heavy and light chains into a single representation, despite their distinct genetic origins and potentially different contributions to antigen recognition (Schatz and Ji, 2011). Full-chain pooling can further dilute binding-relevant complementarity-determining region (CDR) signal with conserved framework sequence, and relatively few studies test whether representations learned from natural complexes transfer to mutation-rich affinity-maturation landscapes (NaderiAlizadeh and Singh, 2025).

Antibody biology suggests a more constrained modelling strategy. Affinity arises from compatibility between a paired antibody and a specific antigen epitope. The antibody heavy chain, antibody light chain and antigen should therefore remain distinguishable until their interaction is modelled. The heavy-chain representation should be enriched for CDR residues, where much of the paratope signal is concentrated, whereas the antigen representation should remain global because the epitope is generally unknown before prediction. Heavy and light chains should exchange context before antibody-level fusion, and antibody-antigen compatibility should be learned through a partner-conditioned mechanism that can suppress protein-language-model features unrelated to binding. For design use, such a model should also provide interpretable residue-level hypotheses, because affinity predictions are more actionable when they can be related to plausible paratope and epitope regions rather than treated as black-box scores (Singh et al., 2021, 2022).

We developed AbAffinity to implement these principles. AbAffinity is a sequence-only, chain-aware three-stream architecture in which heavy-chain, light-chain and antigen sequences are encoded independently using a frozen ESM-2 650M protein language model. The heavy chain is summarised using CDR-focused pooling over CDR-H1, CDR-H2 and CDR-H3, whereas the light chain and antigen are mean-pooled. Heavy-light self-attention and a learned fusion gate produce an adaptive antibody representation, followed by gated cross-attention that models partner-specific compatibility by allowing the antibody representation to selectively filter antigen features (Qiu et al., 2025). A cosine-similarity readout in the shared latent space maps the final antibody and antigen representations to predicted pK_d_ values (Figure 1). Because the protein language model is frozen and only a compact interaction module is trained, AbAffinity remains lightweight for high-throughput sequence scoring. We evaluate AbAffinity across public affinity benchmarks, structure-associated comparisons and affinity-maturation landscapes.

**Figure 1.**
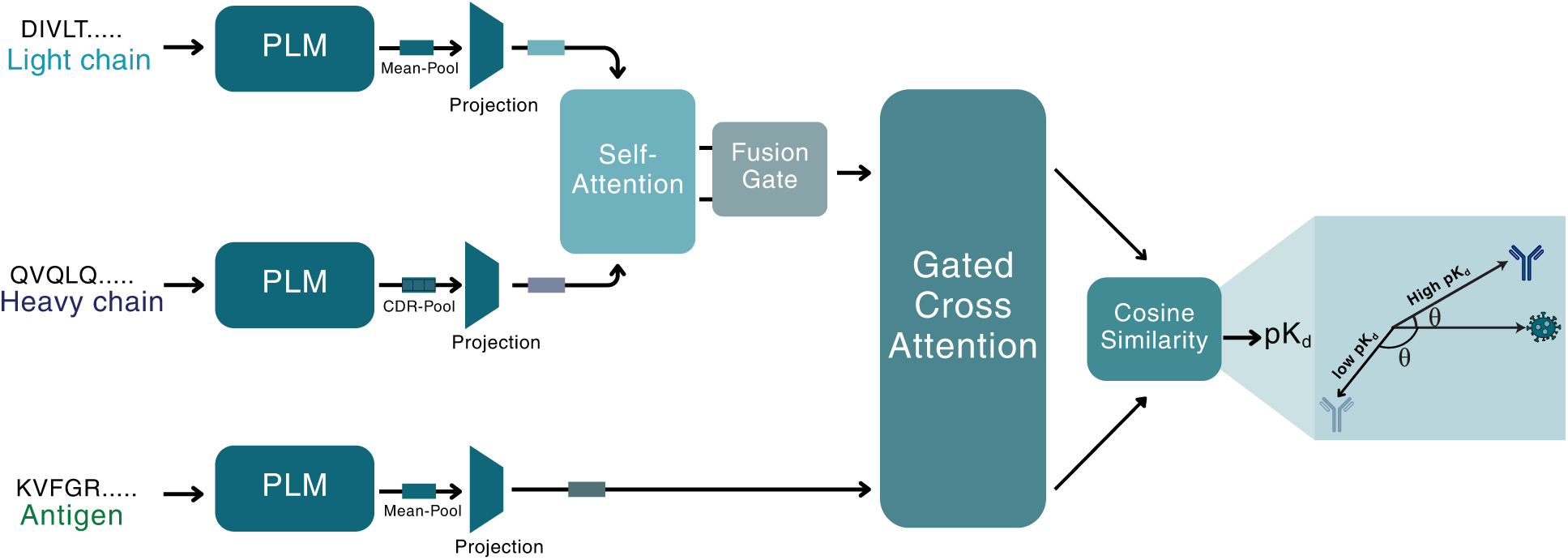
Overview of the AbAffinity framework for antibody-antigen affinity prediction. Heavy-chain, light-chain, and antigen embeddings are projected into a shared interaction space, heavy and light chains exchange information through self-attention, a learned weighted fusion gate forms an antibody representation, and gated cross-attention filters antigen features before separate antibody and antigen heads produce normalized vectors whose cosine similarity is mapped to predicted *pK_d_*.

## 2 Methods

### 2.1 AbAffinity overview

AbAffinity predicts antibody-antigen affinity from antibody heavy-chain, antibody light-chain and antigen sequences. The model has three stages. First, each sequence is independently encoded with a frozen ESM-2 protein language model and pooled into a chain-level representation. Second, heavy-chain, light-chain and antigen representations are projected into a shared latent space, where heavy and light chains are contextualised and fused into an antibody representation. Third, gated antibody-antigen interaction blocks produce an antibody-conditioned antigen representation, and a cosine-similarity readout maps the final antibody and antigen embeddings to predicted pK_d_. Only the downstream interaction network is trained; the ESM-2 encoder remains fixed.

### 2.2 Sequence embeddings and chain pooling

Each antibody heavy chain, light chain and antigen sequence was encoded independently using the frozen ESM-2 protein language model with 650M parameters. For an input sequence *S* of length *L*, the encoder produces residue-level embeddings

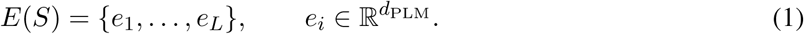

The ESM-2 parameters remain fixed throughout training, and only the downstream projection, interaction and prediction layers are optimised.

To obtain fixed-length chain representations, the heavy chain was summarised using residues assigned to CDR-H1, CDR-H2 and CDR-H3, whereas the light chain and antigen were represented by mean pooling over all residues:

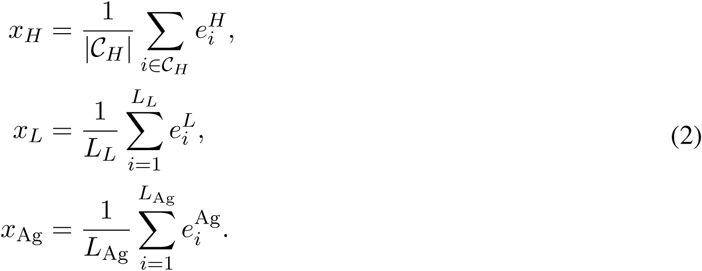

Here, C*_H_* denotes the set of heavy-chain CDR residues, and *L_L_* and *L*_Ag_ denote the light-chain and antigen sequence lengths, respectively.

Heavy-chain CDRs were identified using IMGT numbering (ANARCI) Dunbar and Deane (2016) whenever available. If IMGT numbering failed, a motif-based detector was used to identify CDR-H3. If neither strategy succeeded, the heavy chain defaulted to full-chain mean pooling. This pooling scheme concentrates the antibody representation on likely paratope residues while retaining a global antigen representation, because the epitope is not known before prediction.

### 2.3 Three-stream projection and antibody fusion

The pooled heavy-chain, light-chain and antigen representations were projected into a shared latent interaction space of dimension *d* = 256:

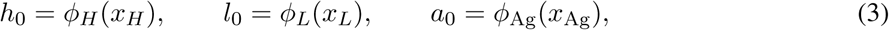

where *ϕ_H_*, *ϕ_L_* and *ϕ*_Ag_ are stream-specific projection networks. Separate projections allow the frozen protein-language-model features to be adapted to the distinct roles of heavy chain, light chain and antigen.

The projected heavy- and light-chain embeddings were treated as a two-token antibody sequence and contextualised using self-attention (Vaswani et al., 2017):

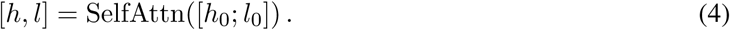

This allows each antibody chain to incorporate information from the other while preserving chain-specific representations.

The contextualised heavy- and light-chain embeddings were then combined through a learned fusion gate:

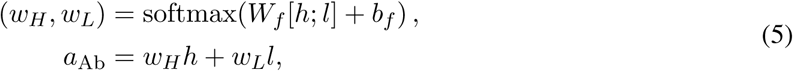

where *w_H_* + *w_L_* = 1. This adaptive fusion allows the model to emphasise the heavy chain, distribute weight across both chains, or accommodate single-domain antibodies by relying primarily on the heavy-chain-like representation.

### 2.4 Gated antibody-antigen interaction

The fused antibody representation interacts with the antigen through gated cross-attention. Given an antibody query *q* and an antigen key/value representation *k*, the gated interaction is defined as

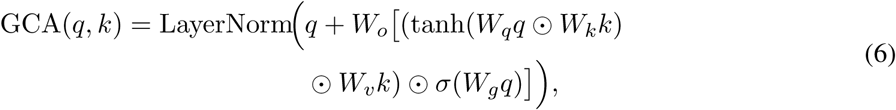

where ⊙ denotes element-wise multiplication and *σ*(·) is the sigmoid activation function. The multiplicative term captures feature-wise antibody-antigen compatibility, while the gate suppresses latent dimensions that are less relevant for the current antibody-antigen pair.

Starting from the projected antigen representation,

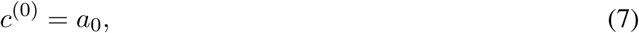

two gated interaction blocks iteratively refine the antigen representation using the fused antibody representation as a fixed query:

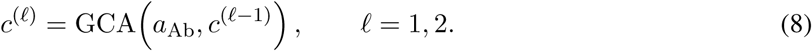

The final output *c* = *c*^(2)^ is used as the antibody-conditioned antigen representation.

### 2.5 Affinity prediction and training objective

The final antibody representation *a*_Ab_ and antibody-conditioned antigen representation *c* were transformed using independent prediction heads and projected onto the unit hypersphere:

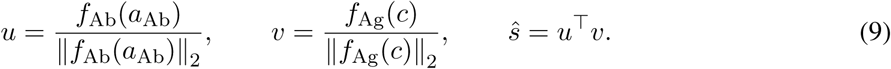

The cosine similarity *s*^ ∈ [−1, 1] serves as the bounded affinity score in the learned interaction space. This bounded readout measures angular alignment between the partner-conditioned antibody and antigen representations, matches the numerical range of the scaled target and limits unconstrained magnitude-based shortcuts that can arise with a concatenation-plus-MLP readout. We interpret this as an architectural inductive bias rather than a direct physical energy model.

During training, experimental pK_d_ values were linearly scaled to [−1, 1] using only the minimum and maximum pK_d_ values observed within the corresponding training fold. The model was optimised using mean-squared error between the predicted similarity *s*^ and the scaled experimental affinity. During inference, every prediction was inverse-transformed to the physical pK_d_ scale using the same training-fold extrema-never the evaluation-fold extrema-before calculating Pearson correlation, Spearman correlation or RMSE:

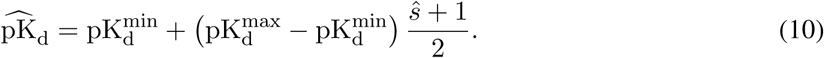

This formulation treats affinity prediction as similarity learning in a shared antibody-antigen embedding space while preserving direct correspondence with experimentally measured affinity values.

### 2.6 Integrated Gradients attribution

To interpret residue-level contributions, the trained model was wrapped with a differentiable pooling layer that allows gradients to propagate from the scalar affinity prediction back to individual residue embeddings. Residue importance was quantified using Integrated Gradients (Sundararajan et al., 2017):

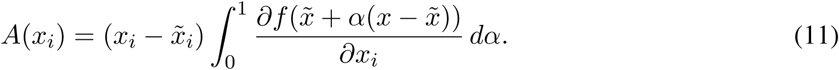

Here, *x_i_* denotes the residue embedding, *x̃_i_* denotes the baseline embedding and *f* denotes the scalar affinity prediction. Attribution scores were summed across embedding dimensions and min-max normalised within each chain for visualisation. The resulting residue-level scores were mapped onto antibody and antigen sequences, and, where structures were available, onto antibody-antigen complexes to compare high-attribution residues with known paratope and epitope regions.

### 2.7 Datasets and experimental setup

The primary training and cross-validation resource was SAAINT-DB, a structurally resolved antibody-antigen interaction database with PDB-linked chain assignments and experimental affinities (Huang et al., 2025). We retained entries with an experimentally measured dissociation constant, an associated PDB ID, antibody heavy chain, antibody light chain, antigen sequence and a finite pK_d_ value. Affinities were converted as pK_d_ = − log_10_(*K_d_*[M]). Across the 2,575 modelling-ready complexes, pK_d_ ranges from 4.333 (approximately 46.5 *µ*M) to 13.469 (approximately 0.034 pM), with mean 8.52. The data comprise 1,913 conventional paired antibodies and 662 single-domain nanobodies/VHHs from 1,271 PDB structures. We evaluated: (i) random ten-fold cross-validation, (ii) a PDB-disjoint split, in which all complexes from the same PDB identifier remain in one fold, and (iii) antigen-cluster-disjoint splits constructed after direct antigen-sequence clustering at 90% and 70% identity. Antigen sequences were clustered using a greedy single-pass procedure: sequences were sorted by length (longest first), and each sequence joined the first existing cluster whose representative was within the identity threshold, or founded a new cluster otherwise. Sequence identity was approximated using the Ratcliff/Obershelp similarity ratio, computed after a length-ratio prefilter. For every outer fold, early stopping and checkpoint selection used an inner validation subset carved exclusively from the outer training data, the outer evaluation fold remained unseen until final evaluation.

External generalisation was assessed using the MVSF-AB release (Li et al., 2024), including natural-complex benchmarks (SAbDab, *n* = 578, held-out Benchmark, *n* = 38) and mutational benchmarks (AB-Bind, *n* = 1,089, SKEMPI 2.0, *n* = 387). Binding free energies in these datasets were harmonised to the pK_d_ scale to maintain consistency across all benchmarks and with the AbAffinity prediction formulation. For affinity-maturation transfer, we used single-antigen variant landscapes from AbBiBench (Zhao et al., 2025) and FLAb2 (Chungyoun and Gray, 2025), scoring variants in zero-shot mode and then fine-tuning with 10-30% labelled data. AAYL51 was reserved for Top-K binder recovery. The detailed curation strategy and all dataset counts are provided in Supplementary Section 7 and Supplementary Table 4.

## 3 Results

### 3.1 AbAffinity outperforms sequence baselines across natural and mutational benchmarks and generalizes to held-out complexes

#### 3.1.1 Within-Dataset Cross-Validation on SAAINT-DB

We first evaluated whether antibody-antigen affinity can be predicted accurately from sequence alone on SAAINT-DB (Huang et al., 2025). SAAINT-DB provides paired antibody, nanobody, antigen and affinity annotations, making it a suitable primary benchmark for testing whether a model can learn affinity-relevant sequence features without using complex structures at inference time. In our processed affinity subset, the dataset contains conventional heavy-light chain antibodies and single-domain nanobodies spanning a broad pK_d_ range, enabling evaluation of both absolute affinity prediction and rank ordering across antibody formats, as shown in Supplementary Figure 8 and detailed in Supplementary Table 4.

Under nested random ten-fold cross-validation, AbAffinity achieved Pearson *r* = 0.838 ± 0.033, Spearman *ρ* = 0.828 ± 0.027 and RMSE = 0.738 ± 0.078 pK_d_, outperforming the fused two-stream and concat+MLP baselines (Table 2). On the PDB-disjoint split, performance decreased to *r* = 0.567 ± 0.018, *ρ* = 0.544 ± 0.014 and RMSE = 1.126 ± 0.027 pK_d_. Since PDB disjointness does not exclude homologous antigens, we additionally clustered antigen sequences at 70% and 90% sequence identity thresholds. step led to a performance drop wherein *r* = 0.449 ± 0.143 with RMSE = 1.244 ± 0.203 pK_d_ at 90% identity and *r* = 0.371 ± 0.135 with RMSE = 1.299 ± 0.161 pK_d_ at 70% identity. These results demonstrate that related-antigen similarity contributes to random-split performance and that extrapolation to sequence-divergent antigens remains a generalisation challenge.

#### 3.1.2 Protocol-Matched Comparisons on External Benchmarks

Beyond SAAINT-DB, AbAffinity was evaluated on established natural and mutational affinity collections. On the natural SAbDab dataset, AbAffinity achieved a Pearson correlation of 0.60 ± 0.02 under ten-fold cross-validation. To evaluate performance on unknown natural antibody-antigen complexes, we trained on SAbDab and evaluated once on the independent 38-complex Benchmark set curated by Guest et al. (2021a). Following the MVSF-AB protocol, this set is treated as sequence-disjoint from the SAbDab training distribution based on the reported CD-HIT analysis. AbAffinity reached Pearson *r* = 0.564 ± 0.026, Spearman *ρ* = 0.570 ± 0.033 and RMSE = 1.364 ± 0.116 pK_d_. Its correlations outperforms the published MVSF-AB Pearson value of 0.467, whereas MVSF-AB has the lower converted RMSE (1.061 pK_d_). On the mutational datasets AB-Bind and SKEMPI 2.0, AbAffinity yields Pearson correlations of 0.78 ± 0.01 and 0.72 ± 0.01, respectively, under the same ten-fold cross-validation. These results indicate that the same sequence-based architecture remains effective when affinity variation arises from mutations.

Predicted-versus-experimental affinity scatter plots provide the corresponding calibration view as demonstrated in Supplementary Figure 9. Dataset-level uncertainty is reported through three complete cross-validation seeds and 1,000 bootstrap resamples of pooled out-of-fold predictions, using the nested-validation checkpoints, the random-split pooled Pearson correlation is 0.836 with a 95% confidence interval of [0.817, 0.854]. We additionally estimated per-prediction epistemic uncertainty on the held-out Benchmark set using the standard deviation across three independently trained model seeds. Ensemble disagreement was positively associated with absolute prediction error (Pearson *r* = 0.32, *p* = 0.05, Spearman *ρ* = 0.28, *p* = 0.09), indicating a modest but informative signal for flagging uncertain candidates rather than a fully calibrated predictive interval.

**Figure 2.**
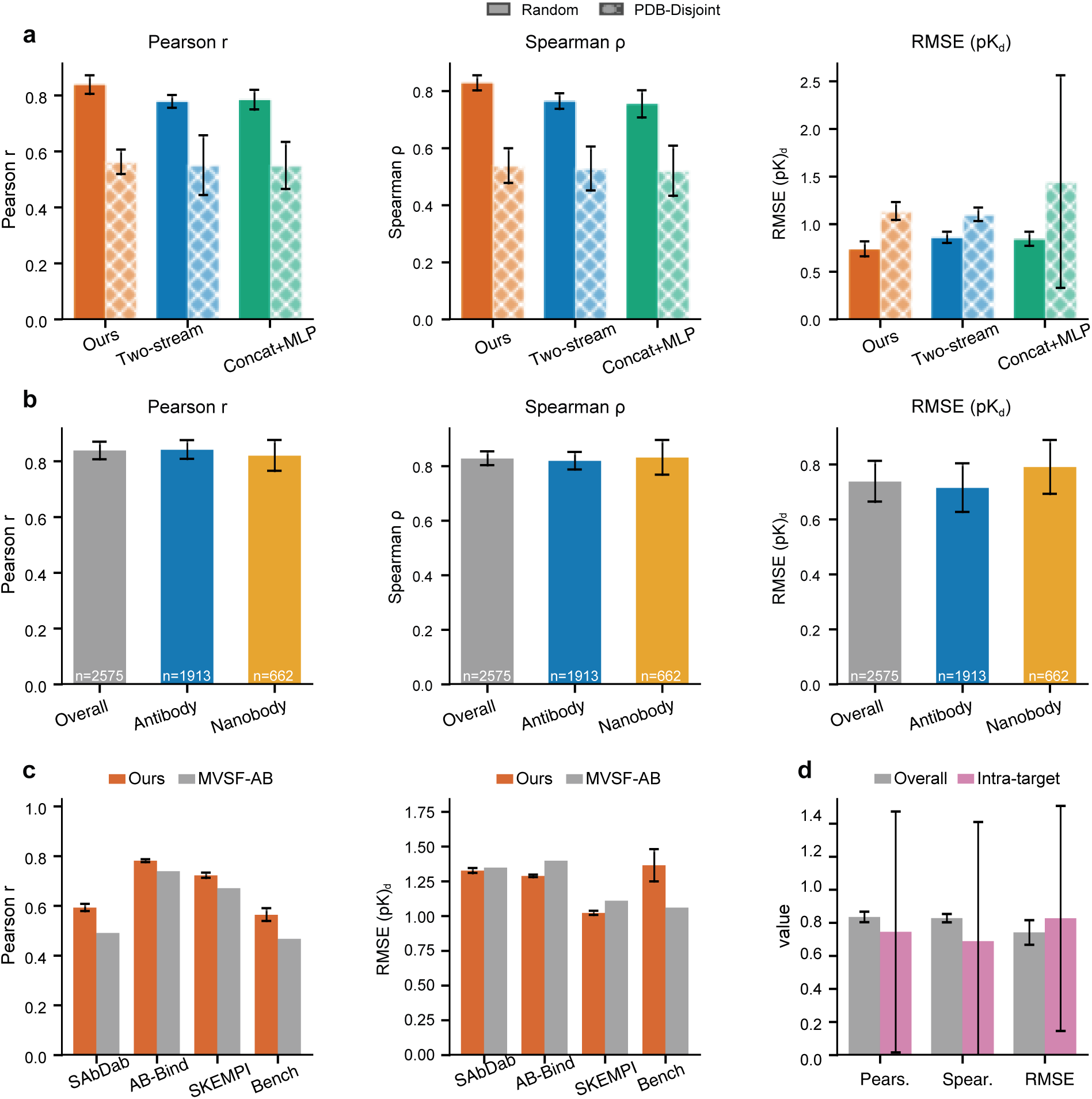
Chain-aware modelling improves affinity prediction across splits and antibody formats. **a,** Architecture comparison between AbAffinity, the fused two-stream baseline and the concat+MLP baseline under nested random and PDB-disjoint splits on SAAINT-DB. **b,** Performance stratified by paired antibodies and nanobodies. **c,** Protocol-matched within-dataset comparisons on SAbDab, AB-Bind and SKEMPI 2.0, together with SAbDab-to-Benchmark transfer. **d,** Overall pooled performance compared with intra-target ranking.

#### 3.1.3 Zero-Shot and Few-Shot Transfer to Mutational Landscapes

We next evaluated whether AbAffinity can support the practical setting of affinity maturation, where the goal is to rank sequence variants against a fixed antigen. For this analysis, we used single-antigen mutational landscapes from AbBiBench and FLAb2, including mutation-rich affinity-optimization tasks designed to evaluate antibody-design models (Zhao et al., 2025). These landscapes differ from the natural-complex setting used for SAAINT-DB training: many variants are closely related to one parental antibody, affinity differences can be local and assay-dependent, and the central task is within-target ranking rather than broad cross-antigen generalisation. This makes them a stringent test of whether the representation learned from natural antibody-antigen complexes can be reused as a prior for mutation-guided design.

**Figure 3.**
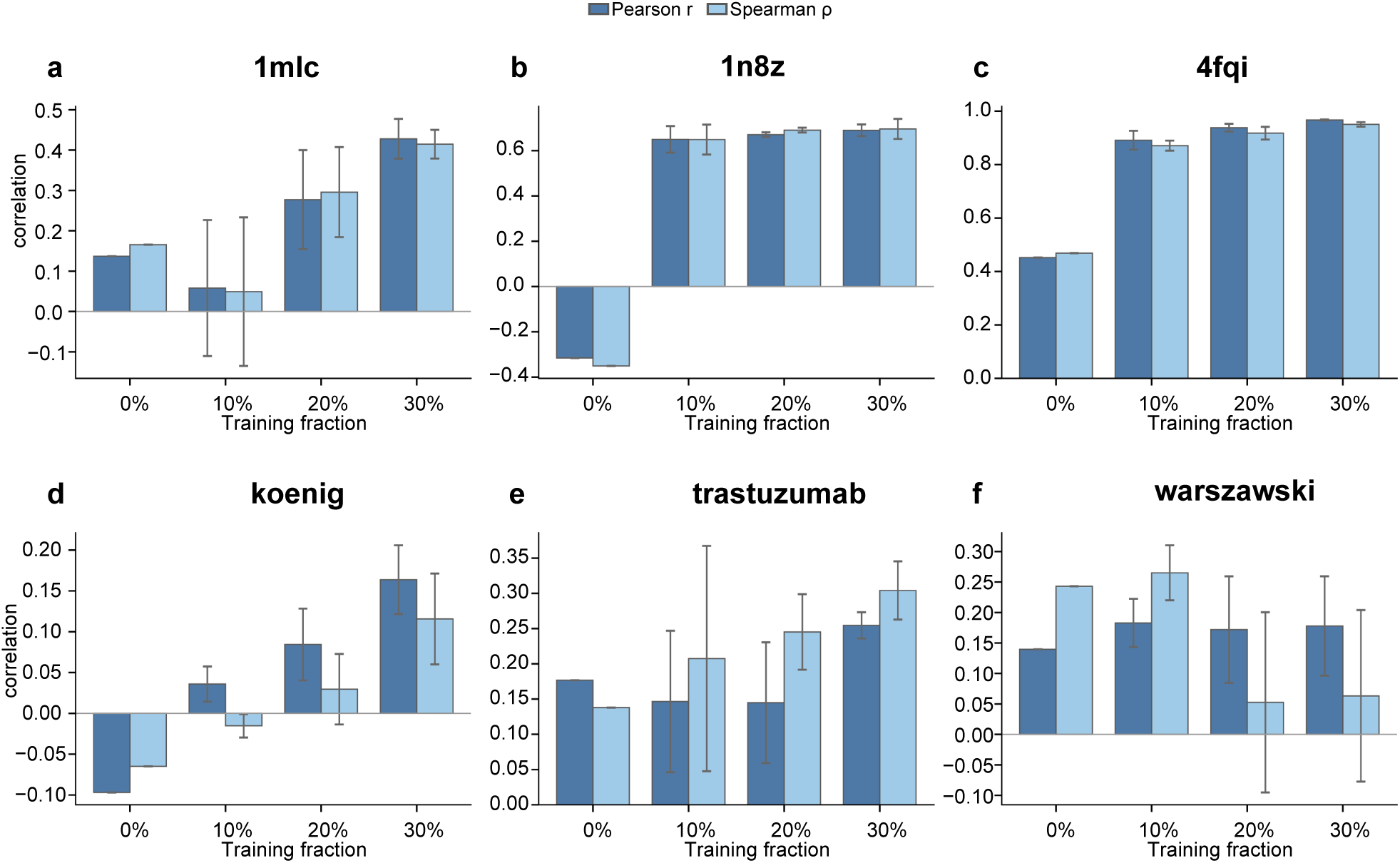
Zero-shot and few-shot transfer to mutational landscapes. Zero-shot transfer and few-shot fine-tuning performance of AbAffinity on six single-antigen mutational landscapes from AbBiBench and FLAb2. Panels (a-c) report Pearson and Spearman correlations after fine-tuning with 10%, 20%, and 30% labelled variants, together with the corresponding zero-shot Pearson correlation. Panels (d-f) evaluate additional affinity-maturation datasets using the same protocol. These results demonstrate that representations learned from natural antibody-antigen complexes can be efficiently adapted to target-specific mutational landscapes, enabling accurate within-antigen ranking with limited labelled data.

In zero-shot mode, AbAffinity retained useful but heterogeneous signal across the three tested landscapes, as shown in Figure 3 and Supplementary Table 20. Zero-shot correlations were positive for 1mlc and 4fqi, with Pearson *r* = 0.14 and *r* = 0.45, respectively, but negative for 1n8z (*r* = −0.32), indicating that direct transfer can fail when the mutational distribution, assay scale or target-specific affinity range differs from the training regime. This behaviour is expected for affinity-maturation datasets, where small local sequence changes can produce target-specific effects that are not fully calibrated by a model trained on diverse natural complexes.

Few-shot fine-tuning substantially improved within-landscape ranking. With only 30% labelled variants, AbAffinity reached Pearson *r* = 0.43 ± 0.05 and Spearman *ρ* = 0.42 ± 0.04 on 1mlc, *r* = 0.69 ± 0.03 and *ρ* = 0.70 ± 0.04 on 1n8z, and *r* = 0.97 ± 0.00 and *ρ* = 0.95 ± 0.01 on 4fqi. The improvement on 1n8z is particularly informative: despite negative zero-shot correlation, limited target-specific calibration converted the pretrained representation into a strong within-target ranker. On 1n8z assay,after fine-tuning with 30% labelled variants, AbAffinity achieved higher correlation than BALM-PPI method, with Pearson *r* = 0.64 ± 0.02 and Spearman *ρ* = 0.60 ± 0.03 under the same labelled-data regime (Singh et al., 2026), as demonstrated in Table 2.

Zero-shot predictions can provide a useful initial prioritisation for some targets, but should be interpreted cautiously when assay conditions or mutation distributions differ from the natural-complex training data. In contrast, few-shot fine-tuning with a modest labelled subset consistently calibrates the model to the local mutational landscape. Thus, AbAffinity acts as a transferable sequence prior: broad antibody-antigen training supplies general binding features, while limited target-specific measurements align the predictions to the antigen, assay and mutation series under optimisation.

### 3.2 Antibody-aware factorisation and CDR-focused pooling explain AbAffinity’s predictive gain

MVSF-AB and AbAffinity both retain heavy-chain, light-chain and antigen inputs, but AbAffinity additionally applies heavy-chain CDR-focused pooling, heavy-light self-attention, gated antibody-antigen cross-attention and a bounded cosine-similarity readout. Under nested ten-fold cross-validation, AbAffinity reached Pearson *r* = 0.838±0.033, compared with 0.784±0.035 for concat+MLP and 0.778±0.023 for the fused two-stream baseline. Because the two-stream baseline retains the interaction and readout design but collapses heavy and light chains before prediction, this comparison supports a contribution from preserving chain identity, it does not imply that three separate input streams alone are unique to AbAffinity.

This chain-aware representation also generalised across antibody formats. AbAffinity handles conventional paired antibodies and single-domain nanobodies without architectural changes: paired antibodies are represented through distinct heavy and light streams, whereas nanobodies are processed by reusing the single-domain chain in the light-chain input position. Across nested-validation checkpoints, the overall Pearson correlation reached 0.838 ± 0.033, paired antibodies reached 0.841 ± 0.033, and nanobodies retained Pearson *r* = 0.820 ± 0.055 and Spearman *ρ* = 0.831 ± 0.064, as shown in Figure 2b and Supplementary Table 17. Thus, the model retains useful performance for both paired antibodies and single-domain binders.

A second biological prior is CDR-focused pooling. Full-chain mean pooling can dilute binding-relevant residues with conserved framework sequence, whereas antibody specificity is concentrated in the complement determining regions. Under the nested-validation protocol, full-chain mean pooling achieved Pearson *r* = 0.809±0.027, Spearman *ρ* = 0.792±0.033 and RMSE = 0.795±0.036 pK_d_. Pooling all heavy-chain CDR residues increased performance to *r* = 0.838 ± 0.033, *ρ* = 0.828 ± 0.027 and RMSE = 0.738 ± 0.078 pK_d_ (Table 1). These analyses indicate that AbAffinity’s gain is not merely a consequence of using a stronger sequence encoder. It arises from imposing antibody-aware structure on the downstream model: heavy and light chains remain identifiable, the heavy-chain representation is concentrated on likely paratope residues, and the antigen is kept separate until partner-specific compatibility is learned. This design also preserves useful within-target ranking. Fisher-aggregated per-antigen correlations remained positive despite larger target-to-target variance, as expected when many antigens contribute only a small number of complexes, as shown in Figure 2d and Supplementary Table 18.

**Table 1.**
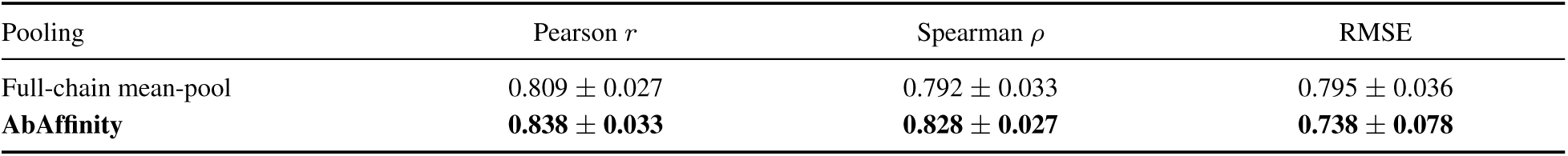
Pooling ablation on SAAINT-DB using nested-validation checkpoints. Values are mean ± s.d., RMSE is reported in pK_d_.

| Table 1. Pooling ablation on SAAINT-DB using nested-validation checkpoints. Values are mean ± s.d., RMSE is reported in $\text{pK}_d$. | | | |
| --- | --- | --- | --- |
| Pooling | Pearson $r$ | Spearman $\rho$ | RMSE |
| Full-chain mean-pool | $0.809 \pm 0.027$ | $0.792 \pm 0.033$ | $0.795 \pm 0.036$ |
| **AbAffinity** | **$0.838 \pm 0.033$** | **$0.828 \pm 0.027$** | **$0.738 \pm 0.078$** |

### 3.3 Gated partner-specific interactions convert protein-language-model features into binding signal

We then tested whether AbAffinity’s performance depends on the upstream protein language model alone, or on the interaction module that converts sequence embeddings into partner-specific binding predictions. Under a matched downstream architecture, ESM-2 provided the strongest frozen sequence representation for antibody-antigen affinity prediction, as demonstrated in Figure 4 and Supplementary Table 9. This encoder comparison used uniform full-chain mean pooling for the non-MINT backbones, rather than the final AbAffinity CDR-focused pooling, so that tokenisation and pooling were treated consistently across encoders. The pooling comparison shows that the final All-CDR configuration reaches *r* = 0.838 ± 0.033, compared with 0.809 ± 0.027 for full-chain mean pooling, indicating that both representation quality and antibody-aware pooling contribute to the final model.

**Figure 4.**
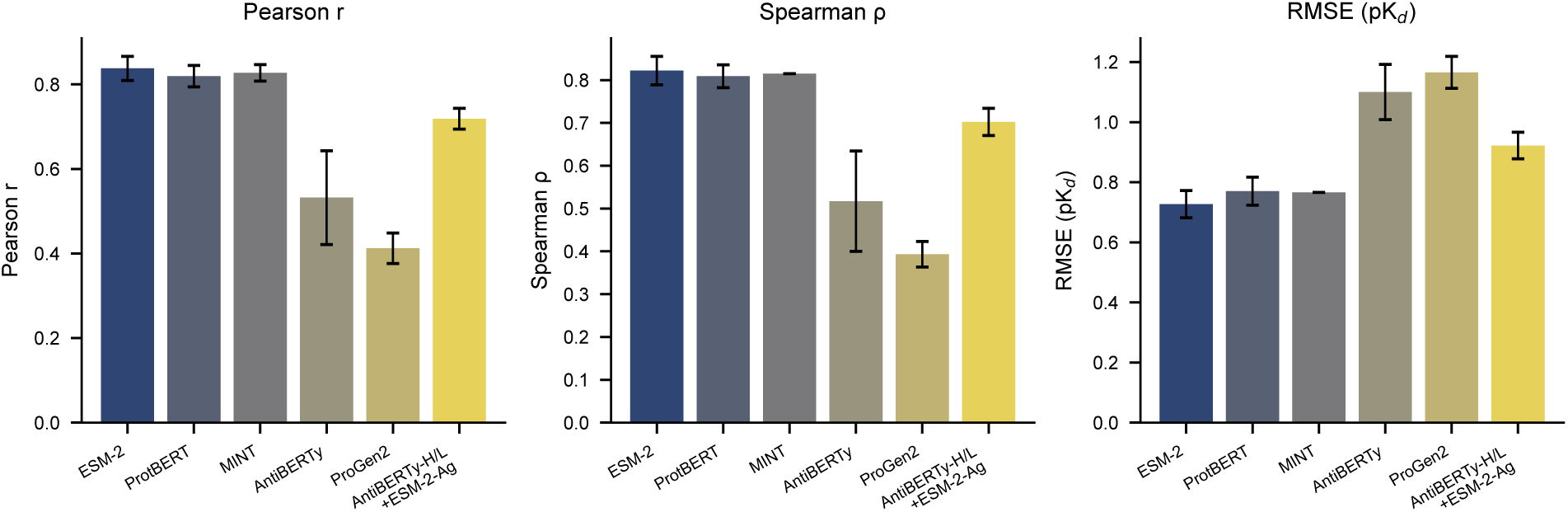
Feature and backbone ablations. Performance comparison of various protein language models-including ESM-2, ProtBERT, AntiBERTy, ProGen2, MINT, and mixed-backbone configurations-as encoders for the downstream AbAffinity architecture. Evaluation metrics include Pearson correlation, Spearman correlation, and RMSE, demonstrating that ESM-2 provides the strongest encoder representation for antibody-antigen affinity prediction.

The encoder comparison also clarifies why a general protein language model is well suited to this task. AntiBERTy is antibody-specialised, but performed poorly when used for both antibody chains in this setting, likely because antigen recognition also requires a general protein representation and because antibody-only pretraining does not encode antigen-side variation. MINT, which is trained to capture protein-protein interaction signal, remained competitive inside the AbAffinity architecture (*r* = 0.83 ± 0.02, *ρ* = 0.82 ± 0.02, RMSE = 0.77 ± 0.04), and was substantially stronger than a frozen MINT+MLP baseline. Nevertheless, it trailed the final ESM-2 configuration. These results suggest that AbAffinity benefits from combining broad evolutionary-scale protein features with an antibody-aware interaction module, rather than relying on any single pretrained encoder alone.

AbAffinity contains two distinct gating mechanisms: the fusion gate, a two-way softmax that forms the antibody representation from the heavy- and light-chain vectors, and the gated cross-attention gate, an independent feature-wise sigmoid gate that modulates the antibody-antigen interaction at each of the two stacked interaction layers. The ablation reported here intervenes only on the latter: gated cross-attention is the main route by which protein-language-model features enter the partner-interaction pathway, and it is the gate this ablation manipulates, applied identically to every gated cross-attention block in the trained model. Re-evaluation of the nested-validation checkpoints gave Pearson *r* = 0.838 ± 0.033 for the learned gate and 0.836 ± 0.036 for a fixed *g* = 0.5 gate, showing that sample-adaptive weighting provides negligible numerical improvement over a uniform mixture for conventional paired antibodies. In contrast, closing the pathway (*g* = 0) reduced correlation to 0.553 ± 0.045 and increased RMSE to 1.124 ± 0.071 pK_d_. Thus, maintaining an active fusion pathway is important, whereas the principal practical benefit of the adaptive gate is architectural flexibility, including accommodation of single-domain antibodies, rather than a large average accuracy gain.

Stream-intervention controls further tested dependence on the supplied partner. Removing the light, heavy or antigen streams reduced Pearson correlation from 0.838 ± 0.033 to 0.764 ± 0.037, 0.592 ± 0.037 and 0.593 ± 0.071, respectively. Replacing the antigen with the dataset mean reduced it to 0.510 ± 0.081, while shuffling antigen partners reduced it to 0.365 ± 0.056 and increased RMSE to 1.330 ± 0.071 pK_d_. The particularly strong effect of shuffling supports dependence on the correct antibody-antigen pairing rather than an antibody-only predictor. These results support dependence on the supplied antigen and its pairing with the antibody. They do not establish fine discrimination among closely related antigens, and input perturbations may also introduce distribution ship

Together, these interventions show that the active interaction pathway and the identity of the supplied antigen materially affect prediction. They support a partner-conditioned compatibility function while leaving open the harder question of discrimination within closely related antigen families. Per-CDR-region mutation analysis further showed that all three heavy-chain CDR loops contribute to prediction, with CDR-H2 best resolved, followed by CDR-H1 and CDR-H3 (Figure 5c and Supplementary Table 19).

**Figure 5.**
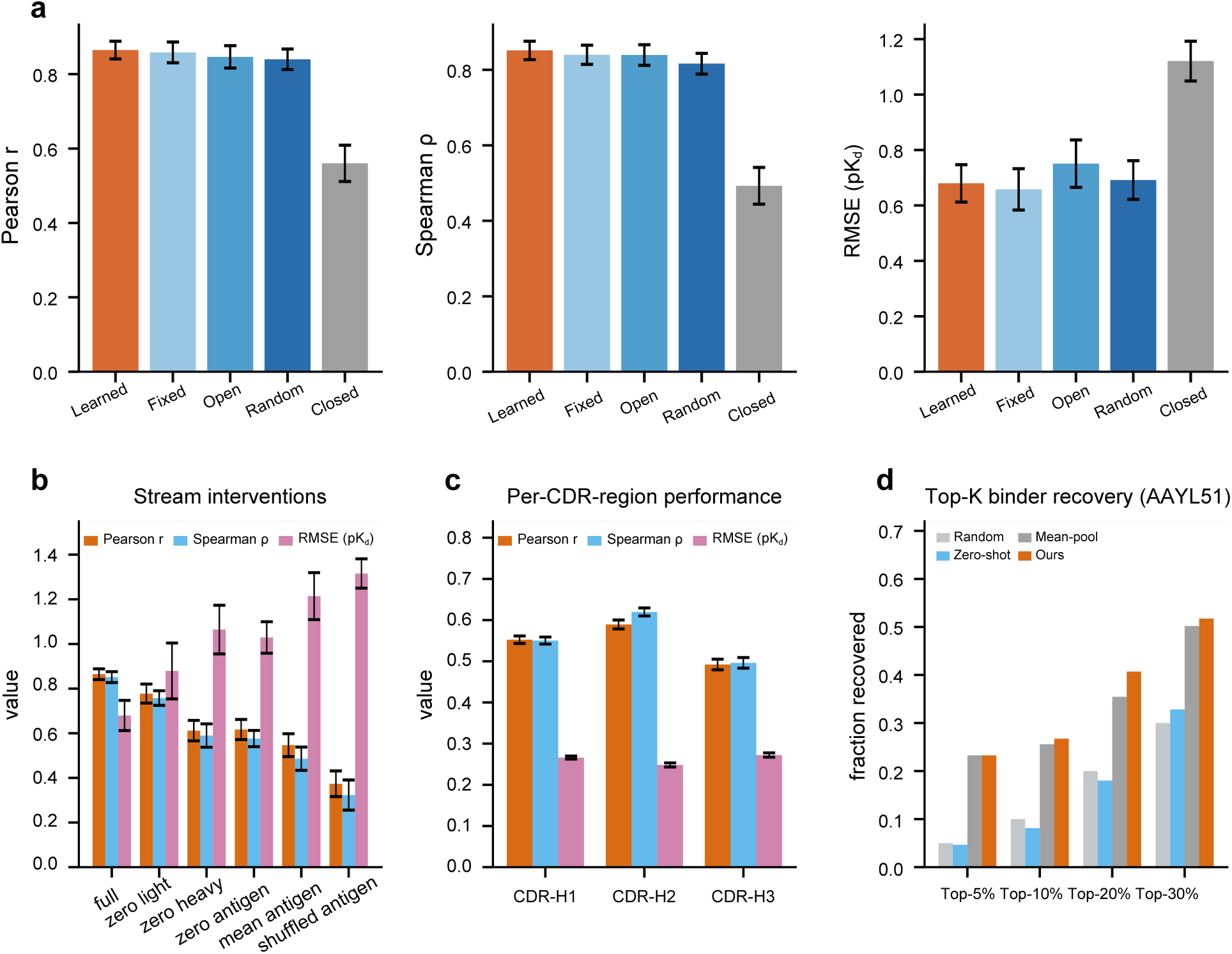
Gating, stream-intervention and mutation-prior controls. **a,** Post-hoc gate interventions comparing learned, fixed, open, random and closed gates. **b,** Stream interventions that remove or corrupt heavy-chain, light-chain and antigen inputs. **c,** Per-CDR-region performance on heavy-chain single mutations. **d,** Top-K binder recovery on AAYL51.

Finally, the same mechanism translated into screening-relevant enrichment. On the AAYL51 mutational landscape, Top-K recovery showed that AbAffinity concentrates experimentally strong binders among its highest-ranked predictions, as demonstrated in Figure 5d and Supplementary Table 21.

At the top-30% cutoff, AbAffinity recovered 51.7% of experimentally top-ranked binders, compared with 30% expected by random selection and 32.8% for zero-shot scoring. It also matched or exceeded the mean-pooling model across cutoffs.

Mutation heatmaps provide design-facing validation beyond global complex-level correlation as presented in Figure 6. The saturation-mutagenesis panel separates permissive CDR positions from affinity-sensitive hot spots, while the substitution matrices show chemically structured effects across residue classes. Conservative substitutions tend to produce smaller changes, whereas charge-altering, polarity-changing and bulky-aromatic substitutions drive larger shifts. These patterns support use of AbAffinity as a compact in silico screen for prioritising CDR positions and substitution classes before experimental testing.

**Figure 6.**
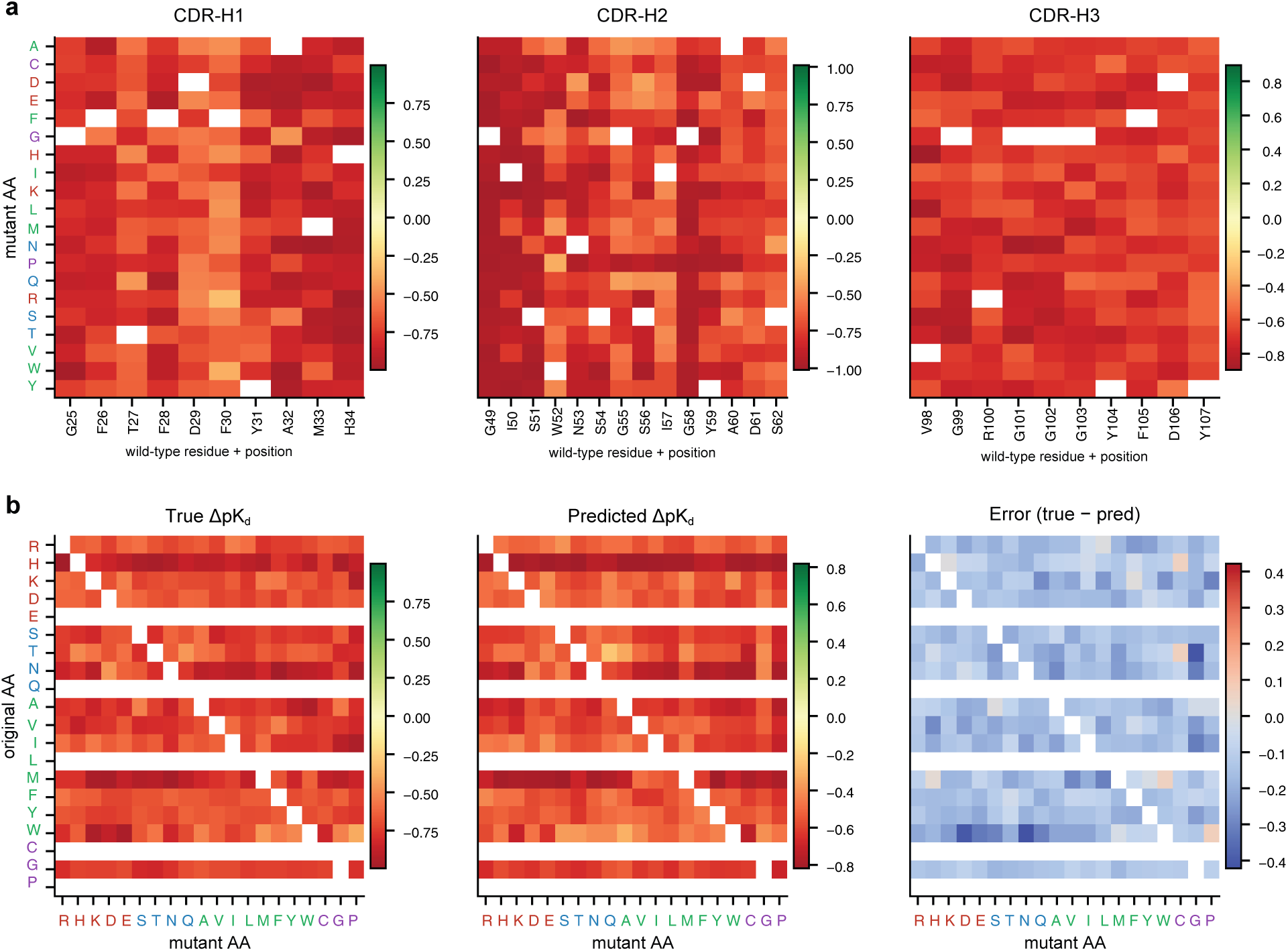
Mutational heatmaps reveal chemically structured affinity effects. **a,** Saturation-mutagenesis heatmap over heavy-chain CDR positions, showing measured mutation-induced changes in affinity. **b,** Wild-type-to-mutant amino-acid substitution matrices comparing measured effects, predicted effects and prediction error, with residues grouped by physicochemical class. This figure tests whether the sequence-only model captures meaningful mutation trends beyond global complex-level ranking, highlighting charged and bulky-aromatic substitutions as major contributors to predicted affinity shifts.

### 3.4 AbAffinity outperforms sequence baselines and remains competitive with structure-based benchmarks

As demonstrated in Table 2, AbAffinity achieves higher Pearson correlations than MVSF-AB on the protocol-matched SAbDab, AB-Bind, SKEMPI 2.0 and held-out Benchmark evaluations. After conversion of all published MVSF-AB errors from kcal/mol to pK_d_, AbAffinity also has lower RMSE on SAbDab, AB-Bind and SKEMPI 2.0. On the 38-complex Benchmark set, however, MVSF-AB has lower absolute error (1.061 versus 1.364 ± 0.116 pK_d_), while AbAffinity has the stronger correlation. We therefore distinguish ranking performance from absolute calibration rather than claiming uniformly lower error.

**Table 2.**
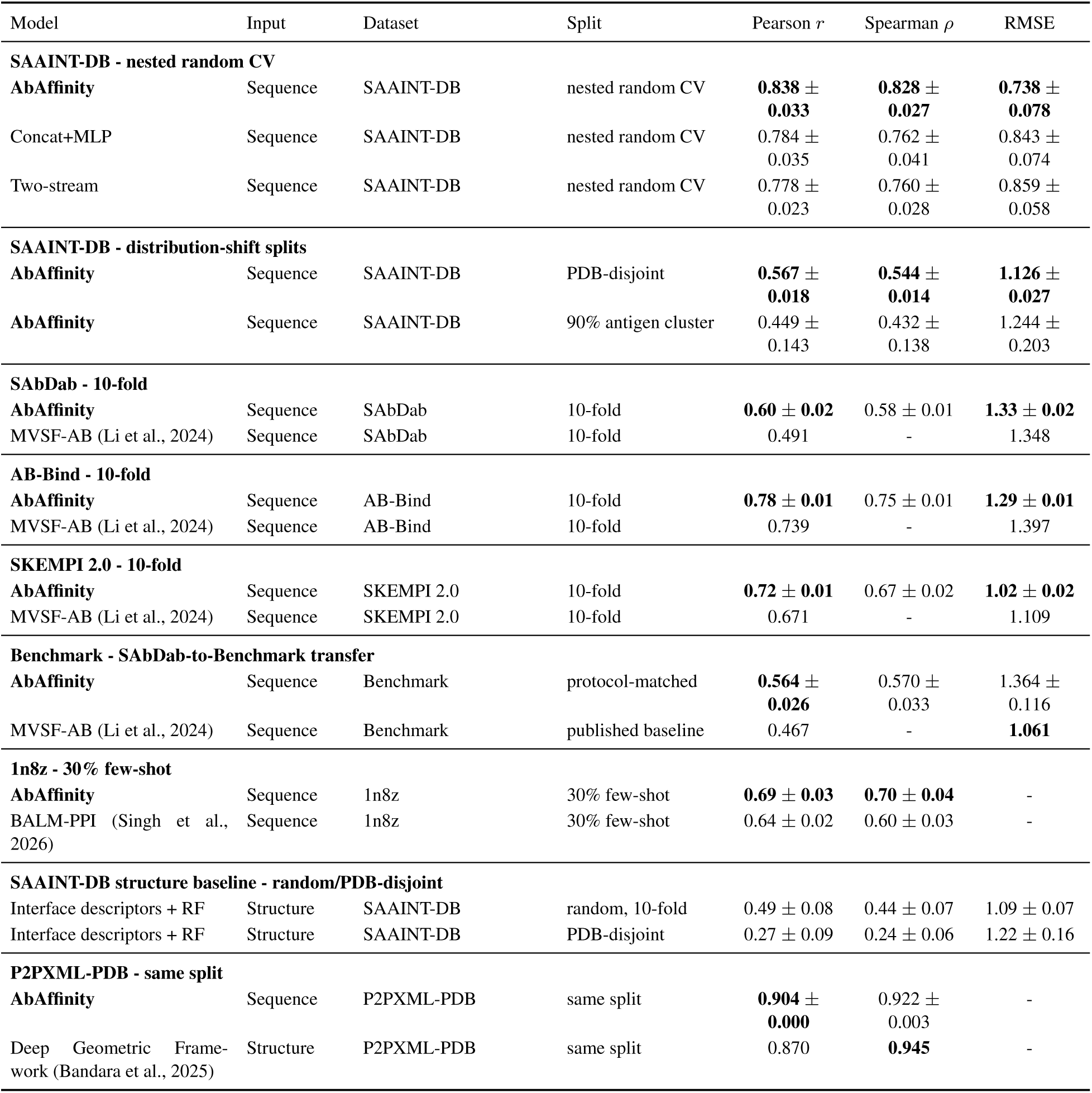
Consolidated benchmark comparison across sequence-only and structure-based affinity methods. SAAINT-DB values use strict nested validation. SAbDab, AB-Bind and SKEMPI 2.0 are within-dataset cross-validation results, whereas Benchmark is SAbDab-to-Benchmark transfer. All RMSE values are in pK_d_ and all AbAffinity inverse transformations use training-set extrema only.

| Model | Input | Dataset | Split | Pearson $r$ | Spearman $\rho$ | RMSE |
| --- | --- | --- | --- | --- | --- | --- |
| **SAAINT-DB - nested random CV** |  |  |  |  |  |  |
| AbAffinity | Sequence | SAAINT-DB | nested random CV | **0.838 ± 0.033** | **0.828 ± 0.027** | **0.738 ± 0.078** |
| Concat+MLP | Sequence | SAAINT-DB | nested random CV | 0.784 ± 0.035 | 0.762 ± 0.041 | 0.843 ± 0.074 |
| Two-stream | Sequence | SAAINT-DB | nested random CV | 0.778 ± 0.023 | 0.760 ± 0.028 | 0.859 ± 0.058 |
| **SAAINT-DB - distribution-shift splits** |  |  |  |  |  |  |
| AbAffinity | Sequence | SAAINT-DB | PDB-disjoint | **0.567 ± 0.018** | **0.544 ± 0.014** | **1.126 ± 0.027** |
| AbAffinity | Sequence | SAAINT-DB | 90% antigen cluster | 0.449 ± 0.143 | 0.432 ± 0.138 | 1.244 ± 0.203 |
| **SAbDab - 10-fold** |  |  |  |  |  |  |
| AbAffinity | Sequence | SAbDab | 10-fold | **0.60 ± 0.02** | 0.58 ± 0.01 | **1.33 ± 0.02** |
| MVSF-AB (Li et al., 2024) | Sequence | SAbDab | 10-fold | 0.491 | - | 1.348 |
| **AB-Bind - 10-fold** |  |  |  |  |  |  |
| AbAffinity | Sequence | AB-Bind | 10-fold | **0.78 ± 0.01** | 0.75 ± 0.01 | **1.29 ± 0.01** |
| MVSF-AB (Li et al., 2024) | Sequence | AB-Bind | 10-fold | 0.739 | - | 1.397 |
| **SKEMPI 2.0 - 10-fold** |  |  |  |  |  |  |
| AbAffinity | Sequence | SKEMPI 2.0 | 10-fold | **0.72 ± 0.01** | 0.67 ± 0.02 | **1.02 ± 0.02** |
| MVSF-AB (Li et al., 2024) | Sequence | SKEMPI 2.0 | 10-fold | 0.671 | - | 1.109 |
| **Benchmark - SAbDab-to-Benchmark transfer** |  |  |  |  |  |  |
| AbAffinity | Sequence | Benchmark | protocol-matched | **0.564 ± 0.026** | 0.570 ± 0.033 | 1.364 ± 0.116 |
| MVSF-AB (Li et al., 2024) | Sequence | Benchmark | published baseline | 0.467 | - | **1.061** |
| **1n8z - 30% few-shot** |  |  |  |  |  |  |
| AbAffinity | Sequence | 1n8z | 30% few-shot | **0.69 ± 0.03** | **0.70 ± 0.04** | - |
| BALM-PPI (Singh et al., 2026) | Sequence | 1n8z | 30% few-shot | 0.64 ± 0.02 | 0.60 ± 0.03 | - |
| **SAAINT-DB structure baseline - random/PDB-disjoint** |  |  |  |  |  |  |
| Interface descriptors + RF | Structure | SAAINT-DB | random, 10-fold | 0.49 ± 0.08 | 0.44 ± 0.07 | 1.09 ± 0.07 |
| Interface descriptors + RF | Structure | SAAINT-DB | PDB-disjoint | 0.27 ± 0.09 | 0.24 ± 0.06 | 1.22 ± 0.16 |
| **P2PXML-PDB - same split** |  |  |  |  |  |  |
| AbAffinity | Sequence | P2PXML-PDB | same split | **0.904 ± 0.000** | 0.922 ± 0.003 | - |
| Deep Geometric Framework (Bandara et al., 2025) | Structure | P2PXML-PDB | same split | 0.870 | **0.945** | - |

The comparison with the Deep Geometric Framework on P2PXML (Bandara et al., 2025) tests whether a sequence-only model can compete on the structure-based method’s benchmark. On the same P2PXML-PDB split and log_10_(IC50) target, AbAffinity reaches Pearson *r* = 0.904 ± 0.000 and Spearman *ρ* = 0.922 ± 0.003, compared with 0.870 and 0.945, respectively, for the Deep Geometric Framework (Supplementary Table 12). Thus, AbAffinity has the higher Pearson correlation, whereas the structure-based comparator retains the higher Spearman correlation.

Per-antigen-family analysis clarifies the scope of the P2PXML benchmark. Among 1,128 recoverable test complexes, 930 are annotated as HIV-1, and no SARS-CoV-2, MERS or influenza entries occur in the P2PXML-PDB structural subset used by the comparator. We also examined the larger P2PXML-Seq table: filtering for unit-harmonised log_10_(IC50) values and complete antigen sequences yielded no usable SARS-CoV-2 rows, no usable influenza rows and only 39 MERS rows against a single antigen. This coverage is insufficient for a robust additional structural comparison. We therefore report the existing sequence-based AAYL51 SARS-CoV-2 and 4fqi influenza evaluations separately and do not present them as structure-based benchmarks.

### 3.5 Attributions recover paratope and epitope residues

Integrated Gradients analysis shows that AbAffinity predictions are supported by biologically meaningful residues rather than diffuse sequence background as illustrated in Figure 7. In the 1VFB D1.3-lysozyme example, the ground-truth structural work identifies a discontinuous lysozyme epitope formed by residues 18-27 and 117-125, which make hydrogen bonds and van der Waals contacts with CDR residues from the antibody. The same study highlights lysozyme Gln121 as a specificity-determining position because substitution at this site disrupts D1.3 recognition (Bhat et al., 1994). The attribution map concentrates signal in the antibody CDR region facing this epitope and in lysozyme positions within the two reported epitope segments, including the 117-125 region that contains Gln121.

**Figure 7.**
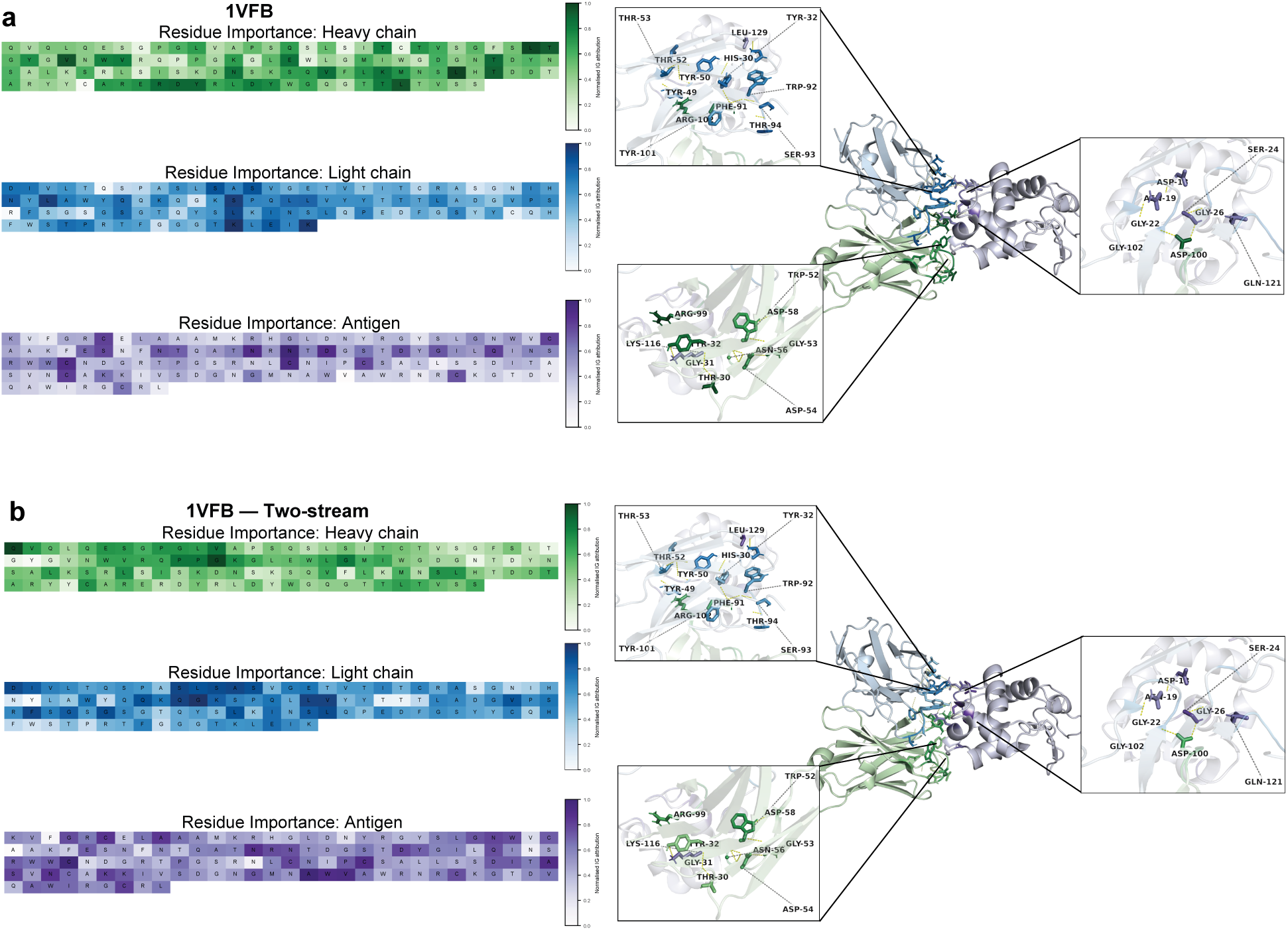
Integrated Gradients attribution and structural mapping for the D1.3-lysozyme complex (PDB: 1VFB). Ground-truth interface residues were referenced from the D1.3-lysozyme crystal-structure study (Bhat et al., 1994). **a,** AbAffinity combines residue-level heatmaps with structural mapping of high-attribution residues onto the antibody-antigen complex. The strongest signals localise to antibody CDR residues and lysozyme epitope positions, showing that the sequence-only model recovers a physically plausible paratope-epitope pattern. **b,** The matched two-stream baseline retains some interface signal but distributes attribution more diffusely across the fused antibody representation. This direct comparison shows why explicit heavy/light factorisation improves explainability: AbAffinity separates chain-specific paratope contributions and maps them more sharply onto known interface residues.

This agreement is important because these residues define the experimentally observed paratope-epitope contact network. At the main-text normalised-attribution threshold of 0.4, AbAffinity recovers 56 of 73 known heavy- and light-chain paratope residues, compared with 22 of 73 for the two-stream baseline. The advantage persists across thresholds: recovery is 64/73 versus 32/73 at *>* 0.3, 47/73 versus 9/73 at *>* 0.5, and 26/73 versus 4/73 at *>* 0.6 (Supplementary Table 22). Integrated Gradients is used here to assess spatial recovery of known interface regions, not as a quantitative energetic or computational alanine-scanning decomposition. The same residue-level analysis is shown for additional complexes in Supplementary Figures 10-12.

## 4 Discussion

AbAffinity shows that sequence-only antibody-antigen affinity prediction can be improved by matching model structure to the biological structure of the binding problem. Rather than treating an antibody-antigen complex as a single concatenated sequence or prematurely fused embedding, AbAffinity preserves the heavy chain, light chain and antigen as separable sources of information until partner compatibility is learned. The resulting gains across SAAINT-DB, external natural-complex benchmarks, mutation-enriched datasets and affinity-maturation landscapes suggest that antibody-aware inductive bias remains valuable even when using strong pretrained protein language models.

A central implication is that protein-language-model embeddings are not sufficient on their own: how they are pooled, factorised and coupled affects affinity prediction. Antigen shuffling shows that the correct pairing matters, but does not by itself establish fine-grained discrimination within closely related antigen families. Likewise, the near-equivalence of learned and fixed gates shows that an active interaction pathway is important while sample-adaptive gate weighting contributes little average numerical gain for conventional paired antibodies. These boundaries temper the mechanistic interpretation while preserving the central architectural result from matched nested-validation baselines.

The transfer experiments clarify the practical role of AbAffinity. In broad screening settings, the model can rank antibody or B-cell receptor candidates directly from sequence, allowing structural modelling and experimental assays to be reserved for a smaller set of prioritised designs. In affinity-maturation settings, zero-shot predictions can provide an initial triage, but few-shot calibration is more reliable when local mutation effects, assay conditions or affinity ranges differ from the training data. Thus, AbAffinity is best viewed as a transferable sequence prior that can be rapidly adapted to a target-specific landscape once a modest number of measurements is available.

The attribution analyses provide an additional check on biological plausibility. Although Integrated Gradients should not be interpreted as a direct energetic decomposition, the concentration of signal on antibody CDRs and known antigen interface regions indicates that the model’s predictions are not driven solely by diffuse sequence background. The sharper recovery of paratope residues relative to the fused two-stream baseline further suggests that explicit chain factorisation improves not only accuracy but also the interpretability of residue-level hypotheses. In practice, these attributions may help nominate CDR positions, antigen-facing regions or mutation classes for experimental review.

Several limitations remain. The reduction from nested random cross-validation (*r* = 0.838) to PDB-disjoint (*r* = 0.567), 90% antigen-cluster-disjoint (*r* = 0.449) and 70% antigen-cluster-disjoint (*r* = 0.371) evaluation shows that related-target information contributes materially to random-split performance. The frozen 650M-parameter ESM-2 backbone and the persistence of the gap after correcting checkpoint selection argue against attributing the entire difference to classical parameter overfitting, nevertheless, extrapolation to new epitope surfaces remains difficult. Zero-shot transfer to mutational landscapes is uneven, and target-specific calibration remains important. Ensemble disagreement provides only a modest uncertainty signal and is not yet a calibrated predictive interval. The model also omits conformational flexibility, induced fit, glycosylation, avidity and assay effects. Finally, Integrated Gradients is model-dependent and should be treated as a spatial hypothesis rather than an energetic decomposition or substitute for alanine scanning.

Future work should therefore move from retrospective prediction toward prospective, uncertainty-aware design. Combining AbAffinity with calibrated uncertainty estimates, selective structural refinement and active learning could enable iterative workflows in which the model proposes informative antibody variants, experiments measure a small batch, and the updated model refines the next design round (Gorantla et al., 2024a; Srivastava et al., 2026). Such closed-loop use would exploit AbAffinity’s main strength: fast, interpretable sequence-based prioritisation before committing to more expensive structural or experimental screening.

## Abbreviations

BCR: B-cell receptor
CDR: complementarity-determining region
IG: Integrated Gradients
PLM: protein language model
RMSE: root mean squared error

## Data availability

The implementation code and source datasets and benchmark releases are openly available from the following repositories and archives:

- AbAffinity implementation: https://github.com/harshitsinghsnu/AbAffinity
- AbAffinity Zenodo archive: DOI: 10.5281/zenodo.23044968
- SAAINT-DB: https://github.com/tommyhuangthu/SAAINT
- MVSF-AB: https://github.com/TAI-Medical-Lab/MVSF-AB
- AbBiBench: https://huggingface.co/datasets/AbBibench/Antibody_Binding_Benchmark_Dataset
- FLAb2: https://github.com/Graylab/FLAb

## 5 Acknowledgements

Harshit Singh would like to acknowledge the support of the Department of Computer Science and Engineering, Shiv Nadar University, Delhi-NCR for providing all the resources needed for computation. Rohan Gorantla acknowledges the support and funding of the United Kingdom Research and Innovation (grant EP/S02431X/1), UKRI Centre for Doctoral Training in Biomedical AI at the School of Informatics, University of Edinburgh, and Exscientia Plc, Oxford.

## 6 Supplementary methods

### 6.1 Sequence embeddings and CDR identification

This subsection builds on ESM-2 protein language model embeddings and antibody CDR annotation conventions (Lin et al., 2023).

Heavy-chain, light-chain and antigen sequences were each encoded with the frozen ESM-2 650M model (`esm2 t33 650M UR50D`), yielding per-residue embeddings of dimension *d*_PLM_ = 1280 from the final representation layer; the terminal `<CLS>`/`<EOS>` tokens were discarded before pooling. To stabilise the scale of the frozen features, each raw chain embedding was layer-normalised over its feature dimension prior to projection. Heavy-chain CDR residues were located with ANARCI under the IMGT scheme, taking CDR-H1, CDR-H2 and CDR-H3 as the IMGT position sets {27-38}, {56-65} and {105-117}, respectively. When ANARCI numbering was unavailable, a motif-based regular-expression detector recovered the three loops from conserved flanking framework patterns (e.g. `C…W[VI]RQ` bounding CDR-H1 and `WYYC…WGQGT` bounding CDR-H3); if neither route resolved a loop, that chain reverted to full-chain mean pooling. The same per-residue caches were reused across all architecture, ablation and seed runs so that the upstream representation remained fixed.

### 6.2 Stream projections and antibody self-attention

This subsection uses the multi-head self-attention formalism introduced for Transformer models (Vaswani et al., 2017).

Each stream-specific projection *ϕ*_{_*_H,L,_*_Ag}_ in Eq. (3) is a block of a linear map to *d* = 256 followed by layer normalisation, a GELU non-linearity and dropout (*p* = 0.1). The heavy/light contextualisation of Eq. (4) is a single multi-head self-attention layer (8 heads, *d* = 256) applied to the length-two token sequence [*h*_0_; *l*_0_], returning the contextualised pair [*h, l*]. The fusion gate of Eq. (5) is a linear map *W_f_* ∈ R^2×2^*^d^* with bias *b_f_* whose softmax yields the convex weights (*w_H_, w_L_*).

### 6.3 Gated cross-attention, expanded

This subsection follows attention-based interaction modelling and gated protein-representation modules (Vasw et al., 2017; Qiu et al., 2025).

The gated cross-attention operator of Eq. (6) is, written with its intermediate terms,

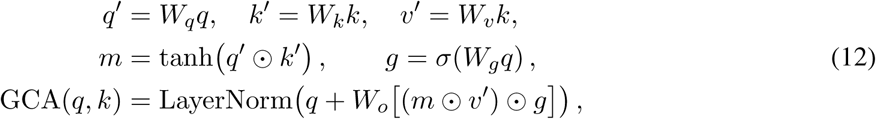

where *W_q_, W_k_, W_v_, W_o_* ∈ R*^d^*^×^*^d^* are bias-free and *W_g_* carries a bias initialised to zero so the gate starts near 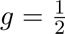. The compatibility term *m* and gate *g* act element-wise over the full *d*-dimensional projected vector. The interaction is stacked for two layers (*n*_layers_ = 2); the antigen context is refined recurrently as in Eq. (8) while the fused antibody query *a*_Ab_ is reused at each layer and as its own output representation. A symmetric antigen→antibody gated block is instantiated in the implementation, but the antibody-conditioned antigen pathway is the route used for the cosine read-out, so the prediction depends on *a*_Ab_ and *c* = *c*^(2)^ only.

### 6.4 Training objective and affinity scaling

For each training fold, experimental affinities were mapped to the cosine range [−1, 1] using only the fold-specific training extrema 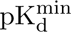 and 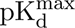,

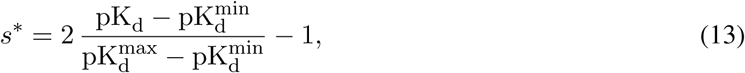

which is the inverse of the inference map in Eq. (10). All reported correlations and errors were calculated only after inverse-transforming predictions with the corresponding training-fold extrema; evaluation-fold labels were never used to set the scale. The bounded target matches the cosine-similarity readout and avoids unconstrained magnitude scaling. The interaction network was trained to minimise the mean-squared error 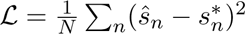 between predicted and target similarities. Only the projection, self-attention, fusion, gated-interaction and read-out parameters were optimised, the ESM-2 backbone remained frozen.

### 6.5 Optimisation and cross-validation protocol

This subsection reports the validation protocol for SAAINT-DB and external MVSF-AB benchmarks (Huang et al., 2025; Li et al., 2024).

Models were optimised with AdamW (learning rate 10^−4^, weight decay 10^−2^), batch size 32, gradient-norm clipping at 1.0 and dropout *p* = 0.1. For every outer evaluation fold, the remaining 90% of the data were divided into an inner training set and an inner validation set. Early stopping and checkpoint selection used only inner-validation Pearson correlation; the outer fold remained unseen until the selected checkpoint was evaluated once. The shared interaction dimension was *d* = 256 with 8 attention heads and two interaction layers. SAAINT-DB was evaluated with nested random ten-fold CV, a PDB-disjoint split, and antigen-cluster-disjoint splits at 90% and 70% sequence identity.

### 6.6 Integrated Gradients implementation

This subsection uses Integrated Gradients for residue-level attribution and maps examples to structurally characterised antibody-antigen complexes (Sundararajan et al., 2017; Bhat et al., 1994; Matho et al., 2012; Liu et al., 2017; Shin et al., 2018).

For attribution, the trained model was wrapped in a differentiable pooling layer that re-derives the pooled chain vectors *x_H_, x_L_, x*_Ag_ from per-residue embeddings, allowing gradients of the scalar prediction *f* to flow back to each residue. Residue importances were computed with Integrated Gradients (Eq. (11)) using a zero-vector baseline *x̃* and 50 Riemann interpolation steps, summed over the *d*_PLM_ embedding dimensions and min-max normalised within each chain to [0, 1] for visualisation. Attributions were computed for AbAffinity and the matched two-stream baseline on the five structurally characterised complexes (1VFB, 4ETQ, 5GRJ, 5Y9J and 3HFM) and mapped onto antibody/antigen sequences and structures.

### 6.7 Affinity conversion for external datasets

This subsection standardises the binding-energy outputs of external MVSF-AB datasets to the pK_d_ scale used for affinity evaluation (Li et al., 2024).

External MVSF-AB datasets report binding free energy Δ*G* (kcal mol^−1^). These were converted to the pK_d_ scale used throughout via

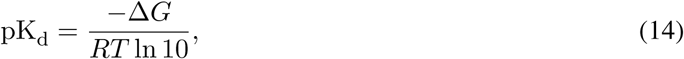

with *R* = 1.987 × 10^−3^ kcal mol^−1^K^−1^ and *T* = 298.15 K, so that all datasets share a single affinity scale for training and evaluation.

### 6.8 Ablations and stream interventions

This subsection compares chain-pooling choices, gated interaction controls and alternative protein language model backbones (Lin et al., 2023; Elnaggar et al., 2021; Ruffolo et al., 2021; Nijkamp et al., 2023; Ullanat et al., 2026).

Pooling ablations compared full-chain averaging with heavy-chain CDR-focused pooling, as demonstrated in Table 1 and Supplementary Table 8. Backbone ablations paired the same downstream architecture with alternative frozen encoders, including ESM-2, ProtBERT, AntiBERTy, ProGen2 and a mixed AntiBERTy antibody plus ESM-2 antigen configuration, as shown in Supplementary Table 9. These analyses tested whether performance depended on the biological pooling prior and on the choice of upstream sequence encoder.

Gate ablations re-evaluated trained models with the gate learned, fixed, fully open, fully closed or randomised while holding all other weights fixed, as shown in Supplementary Table 15. Stream-intervention controls zeroed, averaged or shuffled heavy-chain, light-chain and antigen inputs, as demonstrated in Supplementary Table 16. These controls tested whether predictions depended on the gated partner-aware pathway and authentic antigen information rather than antibody-only regularities.

### 6.9 Transfer and mutation analyses

This subsection evaluates affinity-maturation transfer using AbBiBench and FLAb2-style mutational fitness landscapes (Zhao et al., 2025; Chungyoun and Gray, 2025).

AbAffinity pretrained on natural complexes was applied to three AbBBiBench and three additional FLAb2 landscapes in zero-shot mode and after fine-tuning with 10%, 20% or 30% labelled variants, with AAYL51 evaluated separately for Top-K recovery, as detailed in Supplementary Tables 20 and 21. Fine-tuning was repeated across seeds and evaluated with within-landscape Pearson and Spearman correlations. This setting approximates affinity maturation, where a model trained on broad natural-complex data must adapt to one antigen using limited experimental measurements.

Practical screening utility was measured by Top-K recovery. If *B_K_* is the experimentally top-ranked binder set and *B̂_K_* is AbAffinity’s top-K% predicted set, recovery was computed as

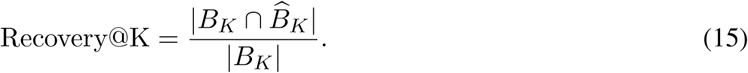

Mutation-level behaviour was further analysed with in-silico saturation-mutagenesis heatmaps over CDR positions and wild-type-to-mutant amino-acid substitution matrices comparing measured effects, predicted effects and prediction error, as shown in Supplementary Tables 19 and 20.

## 7 Supplementary dataset information

The following figure and tables summarise the datasets used for model development and external evaluation, including dataset composition, affinity ranges and filtered benchmark sizes. SAAINT-DB was the only dataset used for primary model training. External MVSF-AB benchmarks were used for cross-dataset evaluation, whereas AbBiBench and FLAb2 (Chungyoun and Gray, 2025) landscapes tested zero-shot and few-shot transfer to affinity-maturation libraries. Dissociation constants were converted to pK_d_ = 9 − log_10_(*K_D_*[nM]); MVSF-AB Δ*G* values were converted using Eq. (14); selection-enrichment assays retained their native fitness scale *Y*.

**Figure 8.**
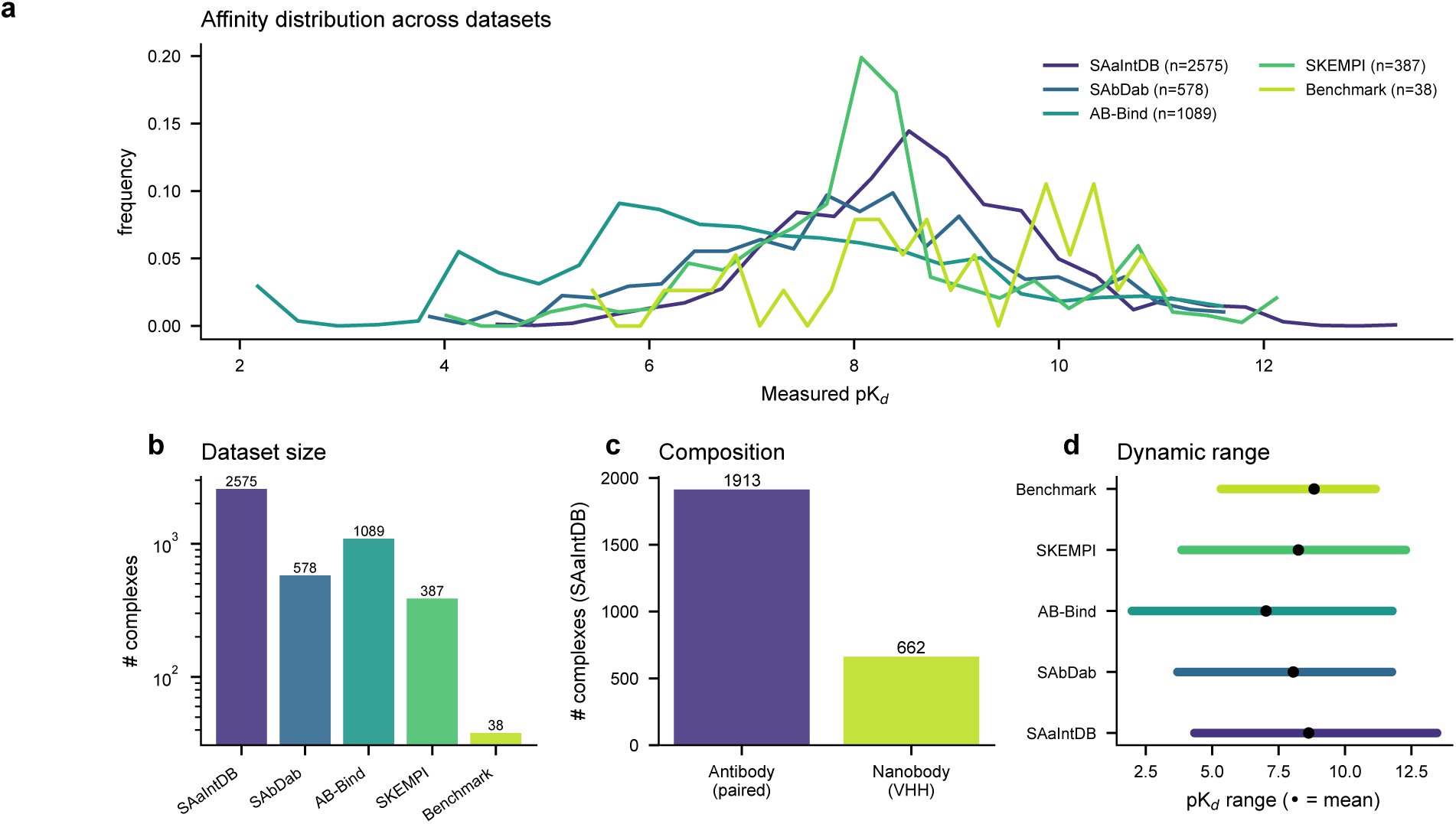
Dataset composition and affinity distributions. **a,** Normalised experimental pK_d_ distributions across SAAINT-DB and the external benchmark datasets, showing overlapping but non-identical affinity regimes. **b,** Dataset sizes on a logarithmic scale, highlighting SAAINT-DB as the primary training resource and the benchmark set as a stringent small held-out test. **c,** Composition of SAAINT-DB, separating paired antibodies from nanobodies. **d,** Dynamic range of pK_d_ values in each dataset, with the mean indicated. This figure defines the prediction problem and shows that the benchmark covers both natural complexes and mutation-driven affinity changes over a broad experimental range.

**Table 3.** Summary of datasets utilized in AbAffinity evaluation, including SAbDab (Dunbar et al., 2014), AB-Bind (Sirin et al., 2016), SKEMPI 2.0 (Jankauskaité et al., 2019) and the processed MVSF-AB release (Li et al., 2024).

| Dataset | Type | Total entries | Available entries |
| --- | --- | --- | --- |
| Kastritis' benchmark | Natural | 144 | 19 |
| Kastritis' benchmark 2.0 | Natural | 179 | 33 |
| SAbDab | Natural | 739 | 579 |
| AB-Bind | Mutant | 1,089 | 1,089 |
| SKEMPI | Mutant | 3,047 | 234 |
| SKEMPI 2.0 | Mutant | 7,085 | 387 |
| Expanded benchmark | Natural | 42 | 38 |

SAAINT-DB curation followed a conservative funnel: 6,158 raw PDB-chain affinity records were filtered to 2,707 entries with measured *K_D_*, collapsed to 2,577 non-redundant complexes, and then reduced to 2,575 modelling-ready examples after excluding two rows with unusable affinity values. The exact affinity range is pK_d_ = 4.333–13.469 (approximately 46.5 *µ*M to 0.034 pM), with mean 8.52. PDB identifiers were retained for the PDB-disjoint split. Direct clustering of antigen sequences produced 601 clusters at 90% identity (largest cluster: 100 complexes); cluster membership, rather than PDB identifier, defined the 90% and 70% antigen-cluster-disjoint folds.

**Table 4.**
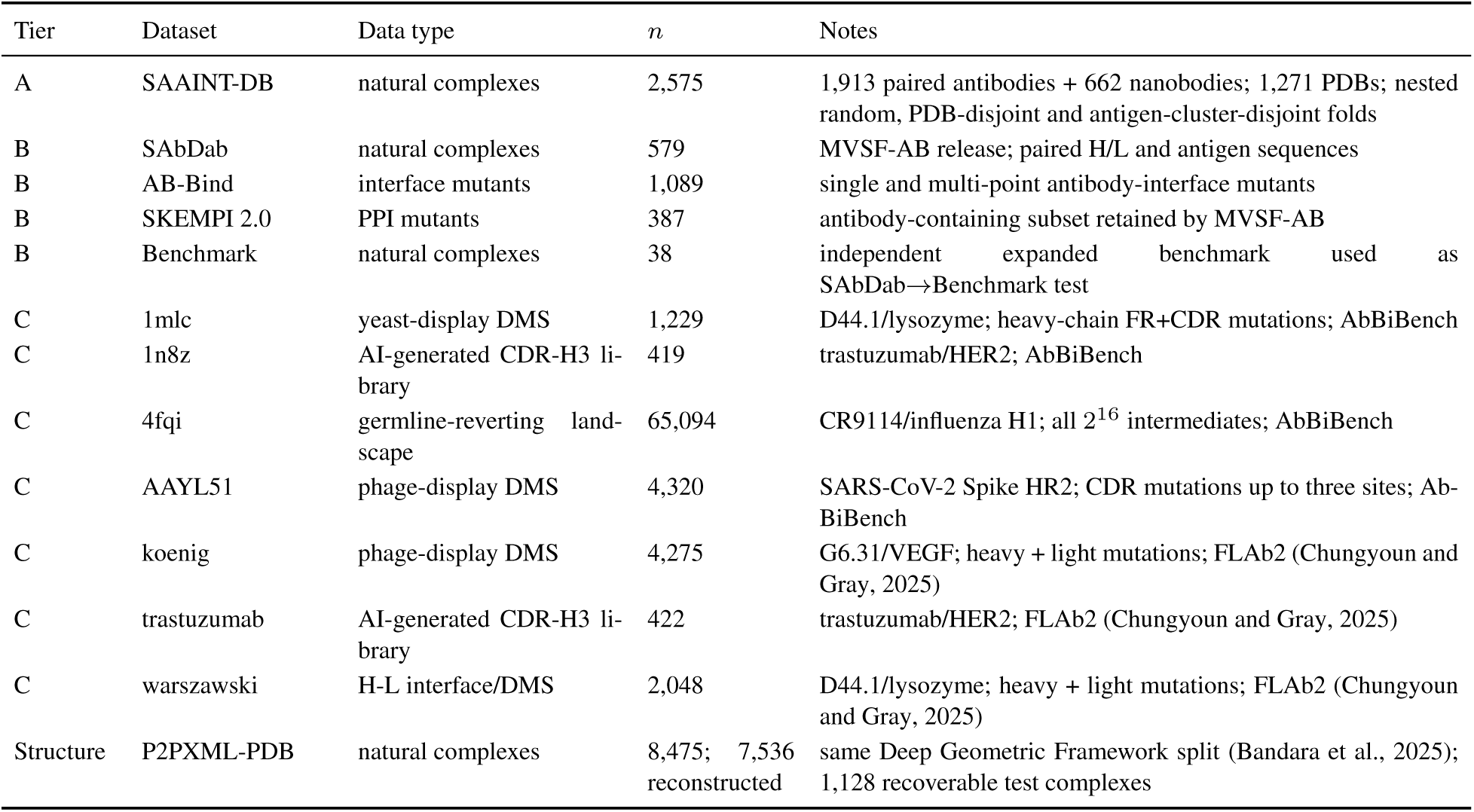
Comprehensive dataset summary. Tier A defines the primary SAAINT-DB training and cross-validation resource; Tier B contains external MVSF-AB benchmarks; Tier C contains single-antigen affinity-maturation landscapes used for zero-shot, few-shot and Top-K transfer analyses.

The MVSF-AB benchmarks were evaluated under their reported protocols to preserve comparability. SAbDab, AB-Bind and SKEMPI 2.0 are within-dataset cross-validation experiments: the model is trained within each dataset and these results are not transfers from SAAINT-DB. The expanded Benchmark set of Guest et al. (2021b) is instead a SAbDab-to-Benchmark transfer evaluation. MVSF-AB reported CD-HIT analysis supporting sequence disjointness between this test set and the SAbDab training distribution. For AbAffinity, cosine outputs were inverse-transformed with SAbDab training-set extrema only. The affinity-maturation landscapes remain separate target-specific ranking tasks: zero-shot scoring uses the SAAINT-DB-trained model directly, few-shot transfer fine-tunes on 10–30% labelled variants with validation-based early stopping, and AAYL51 is analysed by Top-K recovery.

## 8 Supplementary SAAINT-DB validation

These tables report SAAINT-DB held-out validation, pooled uncertainty estimates and architecture comparisons for the SAAINT-DB antibody-antigen affinity resource (Huang et al., 2025). The accompanying scatter plots show predicted versus measured affinity across SAAINT-DB and external datasets.

**Table 5.** SAAINT-DB architecture comparison under strict nested validation. PDB-disjoint and antigen-cluster-disjoint rows quantify increasingly stringent distribution shift.

| Split | Model | Pearson $r$ | Spearman $\rho$ | RMSE |
| --- | --- | --- | --- | --- |
| nested random | **AbAffinity** | **$0.838 \pm 0.033$** | **$0.828 \pm 0.027$** | **$0.738 \pm 0.078$** |
| nested random | Concat+MLP | $0.784 \pm 0.035$ | $0.762 \pm 0.041$ | $0.843 \pm 0.074$ |
| nested random | Two-stream | $0.778 \pm 0.023$ | $0.760 \pm 0.028$ | $0.859 \pm 0.058$ |
| PDB-disjoint | **AbAffinity** | **$0.567 \pm 0.018$** | **$0.544 \pm 0.014$** | **$1.126 \pm 0.027$** |
| 90% antigen cluster | AbAffinity | $0.449 \pm 0.143$ | $0.432 \pm 0.138$ | $1.244 \pm 0.203$ |
| 70% antigen cluster | AbAffinity | $0.371 \pm 0.135$ | $0.327 \pm 0.146$ | $1.299 \pm 0.161$ |

**Table 6.** Per-prediction ensemble uncertainty on the held-out Benchmark set (*n* = 38). Uncertainty is the standard deviation across three independently trained seeds; error is absolute deviation from the experimental pK_d_.

| Association | Correlation | $p$ value |
| --- | --- | --- |
| Pearson: ensemble s.d. versus absolute error | 0.32 | 0.05 |
| Spearman: ensemble s.d. versus absolute error | 0.28 | 0.09 |

**Table 7.**
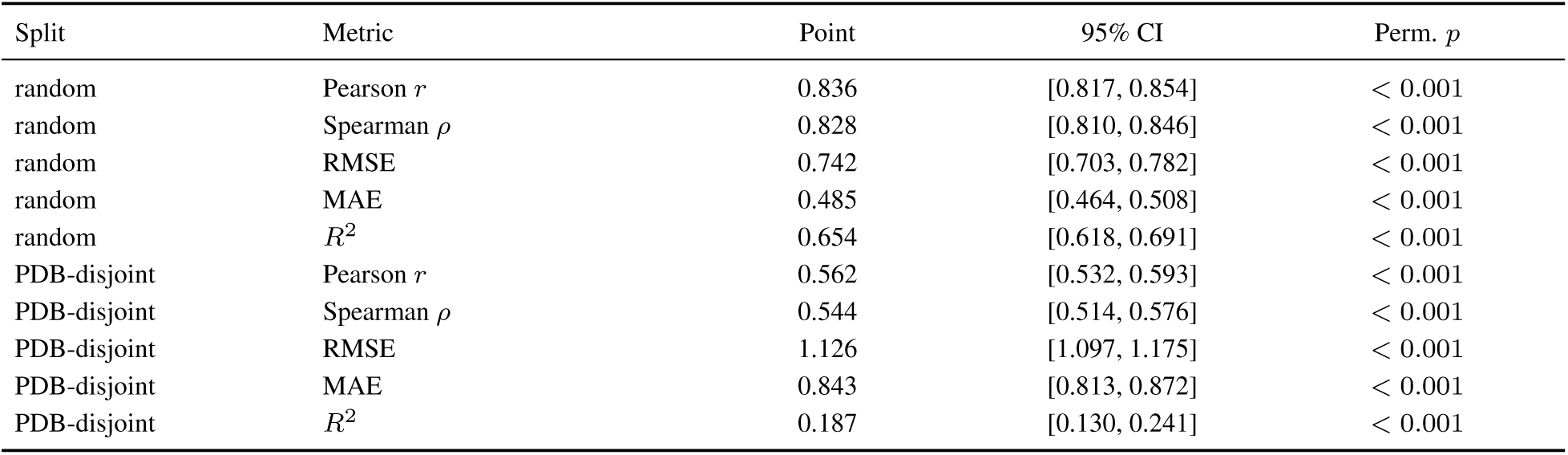
Pooled out-of-fold validation of AbAffinity on SAAINT-DB using corrected nested-validation checkpoints. Point estimates are computed after concatenating one held-out prediction per complex (*n* = 2, 575), so they are dataset-level pooled metrics rather than fold or seed averages. Confidence intervals use 1,000 bootstrap resamples; permutation *p* values use 1,000 label permutations. The PDB-disjoint split corresponds to the split labelled “cold” in the raw evaluation outputs.

| Split | Metric | Point | 95% CI | Perm. $p$ |
| --- | --- | --- | --- | --- |
| random | Pearson $r$ | 0.836 | [0.817, 0.854] | $< 0.001$ |
| random | Spearman $\rho$ | 0.828 | [0.810, 0.846] | $< 0.001$ |
| random | RMSE | 0.742 | [0.703, 0.782] | $< 0.001$ |
| random | MAE | 0.485 | [0.464, 0.508] | $< 0.001$ |
| random | $R^2$ | 0.654 | [0.618, 0.691] | $< 0.001$ |
| PDB-disjoint | Pearson $r$ | 0.562 | [0.532, 0.593] | $< 0.001$ |
| PDB-disjoint | Spearman $\rho$ | 0.544 | [0.514, 0.576] | $< 0.001$ |
| PDB-disjoint | RMSE | 1.126 | [1.097, 1.175] | $< 0.001$ |
| PDB-disjoint | MAE | 0.843 | [0.813, 0.872] | $< 0.001$ |
| PDB-disjoint | $R^2$ | 0.187 | [0.130, 0.241] | $< 0.001$ |

**Figure 9.**
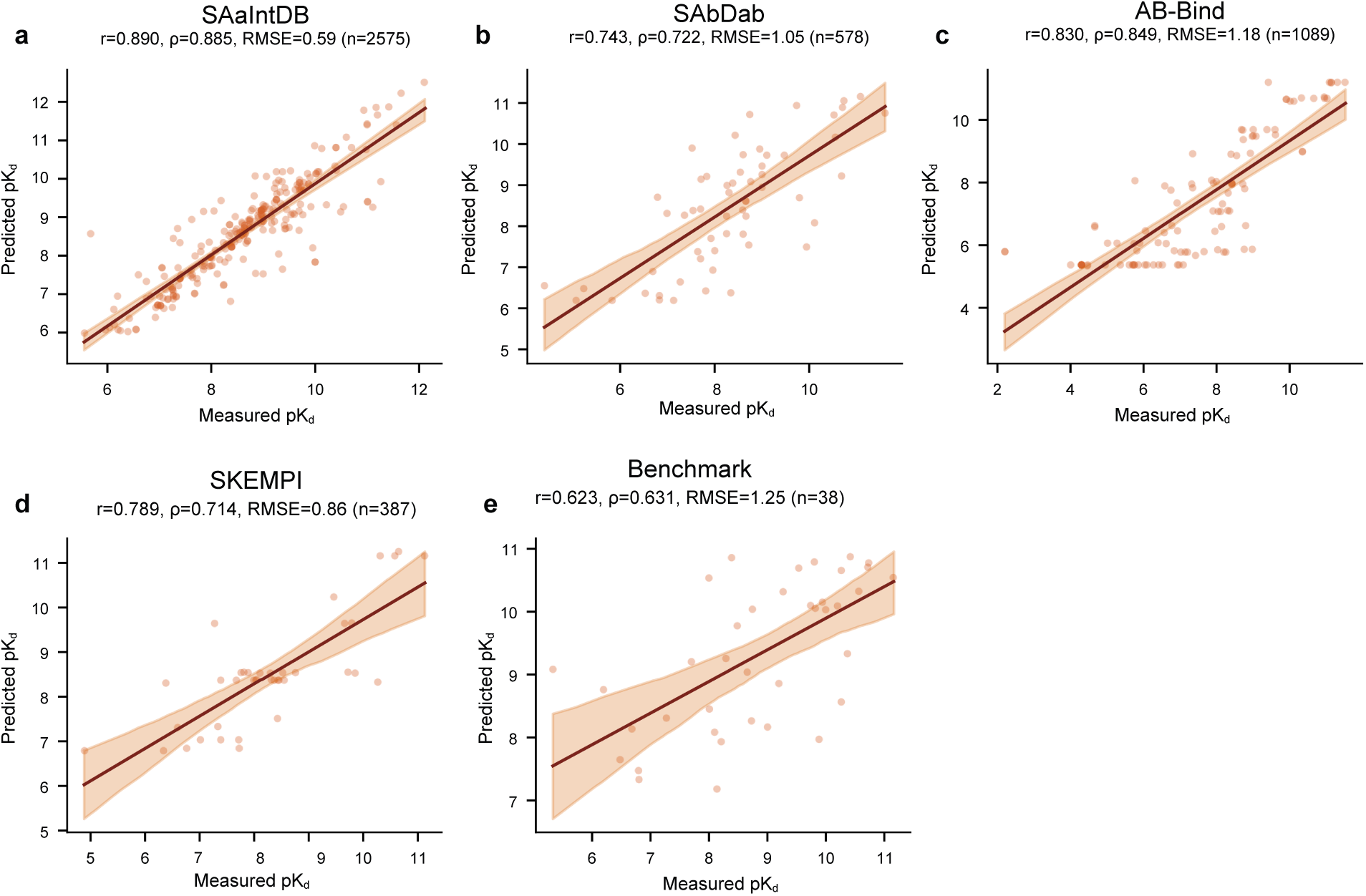
Predicted versus measured affinity across benchmark datasets. Representative out-of-fold scatter plots compare predicted and experimental pK_d_ for SAAINT-DB, SAbDab, AB-Bind, SKEMPI and the held-out benchmark, with ordinary-least-squares fits, bootstrap 95% confidence bands, Pearson *r*, Spearman *ρ*, RMSE and sample size annotated. The positive slopes across all five datasets show that AbAffinity preserves affinity ordering across natural-complex and mutational regimes; the wider bands on smaller sets reflect sample-size limitations.

## 9 Supplementary feature and architecture ablations

These controls isolate the effects of CDR-focused pooling and frozen protein-language-model backbone choice under matched downstream architectures, comparing ESM-2, ProtBERT, AntiBERTy, ProGen2 and MINT representations (Lin et al., 2023; Elnaggar et al., 2021; Ruffolo et al., 2021; Nijkamp et al., 2023; Ullanat et al., 2026).

**Table 8.** Pooling ablation on SAAINT-DB using corrected nested-validation checkpoints. Values are mean ± s.d.; RMSE is reported in pK_d_.

| Pooling | Pearson $r$ | Spearman $\rho$ | RMSE |
| --- | --- | --- | --- |
| Full-chain mean-pool | 0.809 ± 0.027 | 0.792 ± 0.033 | 0.795 ± 0.036 |
| **AbAffinity** | **0.838 ± 0.033** | **0.828 ± 0.027** | **0.738 ± 0.078** |

**Table 9.** Protein-language-model backbone comparison on SAAINT-DB under ten-fold cross-validation, comparing ESM-2 (Lin et al., 2023), ProtBERT/ProtTrans (Elnaggar et al., 2021), MINT (Ullanat et al., 2026), AntiBERTy (Ruffolo et al., 2021) and ProGen2 (Nijkamp et al., 2023). This encoder ablation uses uniform full-chain mean pooling for the non-MINT backbones and one seed; values are mean ± s.d. across the ten folds. Thus the ESM-2 row is the mean-pooled ESM-2 configuration, not AbAffinity.

| PLM | Pearson $r$ | Spearman $\rho$ | RMSE |
| --- | --- | --- | --- |
| **ESM-2 (650M)** | **0.838 ± 0.028** | **0.822 ± 0.033** | **0.727 ± 0.045** |
| ProtBERT | 0.819 ± 0.025 | 0.809 ± 0.027 | 0.770 ± 0.047 |
| MINT (interaction PLM)<sup>‡</sup> | 0.827 ± 0.020 | 0.815 ± 0.024 | 0.766 ± 0.036 |
| AntiBERTy | 0.532 ± 0.111 | 0.517 ± 0.117 | 1.100 ± 0.092 |
| ProGen2 | 0.412 ± 0.036 | 0.393 ± 0.030 | 1.165 ± 0.053 |
| AntiBERTy (H/L) + ESM-2 (Ag) | 0.719 ± 0.025 | 0.702 ± 0.032 | 0.922 ± 0.044 |

<sup>‡</sup>MINT was used as the backbone inside our architecture: per-residue MINT embeddings were pooled over heavy-chain CDR positions and passed through the same gated tri-stream model.

## 10 Supplementary external benchmark comparisons

Unless otherwise stated, benchmark values are reported from matched re-implementations or the corresponding publications, and RMSE is expressed in pK_d_. Corrected nested-validation values are used for AbAffinity, concat+MLP and the two-stream random-split comparison. PDB-disjoint values replace the former “cold” terminology. Dashes indicate metrics not available from the corrected rerun.

**Table 10.** SAAINT-DB comparison under nested random and PDB-disjoint evaluation.

| Split | Model | Pearson $r$ | Spearman $\rho$ | RMSE |
| --- | --- | --- | --- | --- |
| nested random | **AbAffinity** | **0.838 ± 0.033** | **0.828 ± 0.027** | **0.738 ± 0.078** |
| nested random | Concat+MLP | 0.784 ± 0.035 | 0.762 ± 0.041 | 0.843 ± 0.074 |
| nested random | Two-stream | 0.778 ± 0.023 | 0.760 ± 0.028 | 0.859 ± 0.058 |
| PDB-disjoint | **AbAffinity** | **0.567 ± 0.018** | **0.544 ± 0.014** | **1.126 ± 0.027** |
| PDB-disjoint | Concat+MLP | 0.549 ± 0.084 | 0.520 ± 0.088 | 1.445 ± 1.116 |
| PDB-disjoint | DuaDeep-SeqAffinity (reimpl.) | 0.458 ± 0.101 | 0.413 ± 0.108 | 1.238 ± 0.166 |
| PDB-disjoint | DG-Affinity (reimpl.) | 0.436 ± 0.135 | 0.412 ± 0.159 | 1.160 ± 0.091 |

**Table 11.** Protocol-matched comparison with MVSF-AB (Li et al., 2024). SAbDab, AB-Bind and SKEMPI 2.0 are within-dataset ten-fold cross-validation experiments; Benchmark is SAbDab-to-Benchmark transfer. Published MVSF-AB RMSE values were converted from kcal/mol to pK_d_ using *RT* ln 10 = 1.364 kcal/mol at 298.15 K.

| Dataset | AbAffinity Pearson | AbAffinity Spearman | AbAffinity RMSE | MVSF Pearson | MVSF RMSE |
| --- | --- | --- | --- | --- | --- |
| SAbDab | **0.60 ± 0.02** | 0.58 ± 0.01 | **1.33 ± 0.02** | 0.491 | 1.348 |
| AB-Bind | **0.78 ± 0.01** | 0.75 ± 0.01 | **1.29 ± 0.01** | 0.739 | 1.397 |
| SKEMPI 2.0 | **0.72 ± 0.01** | 0.67 ± 0.02 | **1.02 ± 0.02** | 0.671 | 1.109 |
| Benchmark | **0.564 ± 0.026** | 0.570 ± 0.033 | 1.364 ± 0.116 | 0.467 | **1.061** |

## 11 Supplementary P2PXML structural benchmark

These tables place the sequence-only model against the Deep Geometric Framework on the recoverable P2PXML-PDB subset, including full-data, data-efficiency and antigen-family analyses (Bandara et al., 2025).

**Table 12.** Comparison with the Deep Geometric Framework on the P2PXML-PDB benchmark (Bandara et al., 2025). AbAffinity is sequence-only and uses the unchanged tri-stream architecture with cosine readout. The structure-based comparator is the Deep Geometric Framework evaluated on its P2PXML-PDB benchmark with the same log_10_(IC50) target and the same split for fair comparison.

| Model | Pearson $r$ | Spearman $\rho$ |
| --- | --- | --- |
| **AbAffinity (sequence-only)** | **0.904 ± 0.000** | 0.922 ± 0.003 |
| Deep Geometric Framework (structure-based) | 0.870 | **0.945** |

Sequences were reconstructed for 7,536/8,475 P2PXML-PDB complexes (89%) by joining to P2PXML-Seq with PDB extraction as fallback; evaluation uses 1,128 recoverable test complexes. The remaining Spearman gap is consistent with structure retaining an advantage for fine rank ordering among near-equal-affinity complexes.

**Table 13.**
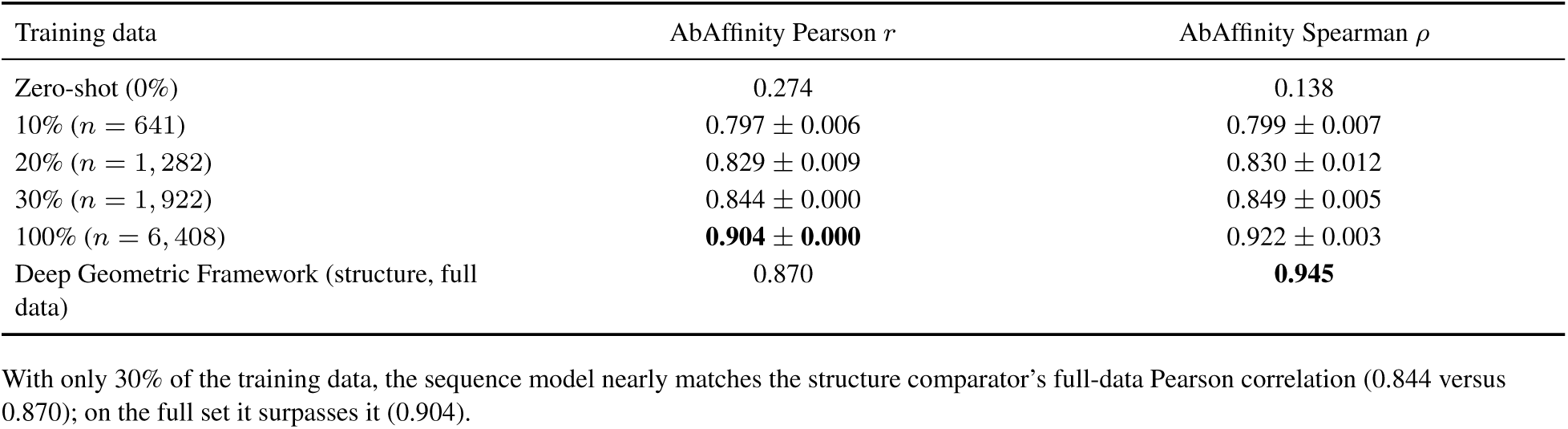
Data efficiency on P2PXML-PDB relative to the Deep Geometric Framework (Bandara et al., 2025). AbAffinity is fine-tuned on fractions of the Deep Geometric Framework training set and evaluated on the same test split as Supplementary Table 12. Zero-shot is a single deterministic frozen-model prediction, and the Deep Geometric Framework full-data values are point estimates from its publication.

**Table 14.**
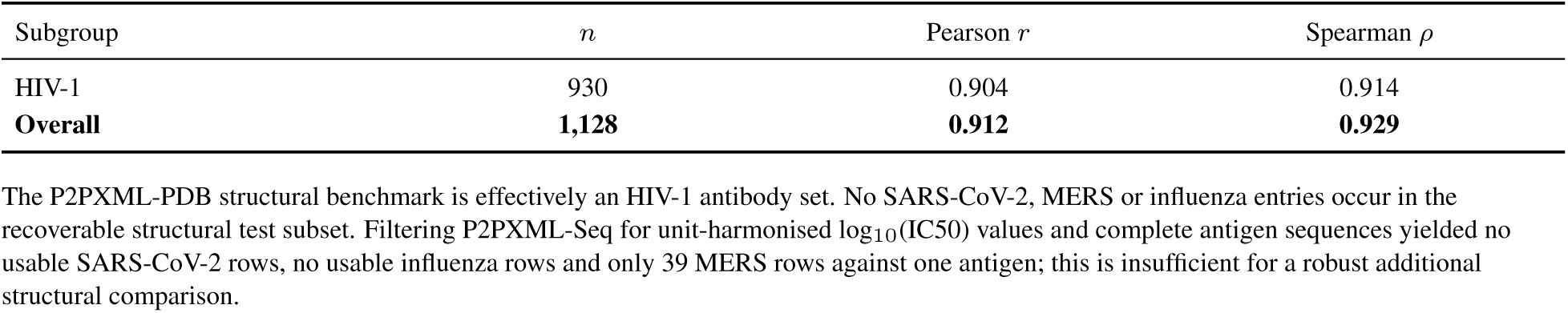
Per-antigen-family breakdown on the recoverable P2PXML-PDB test set. Overall/HIV-1 correlations are computed on the seed-ensemble-averaged prediction

## 12 Supplementary mechanism and stratification analyses

The following tables report gate interventions, stream interventions, antibody-format stratification and intra-target ranking analyses, corresponding to the mechanistic controls described in the Results and to antibody/nanobody benchmark stratification in SAAINT-DB (Huang et al., 2025).

**Table 15.** Gating ablation on SAAINT-DB using corrected nested-validation checkpoints. Post-hoc interventions are evaluated on true outer held-out folds; values are mean ± s.d. across the ten folds.

| Gate | Pearson $r$ | Spearman $\rho$ | RMSE |
| --- | --- | --- | --- |
| Learned (final) | 0.838 ± 0.033 | 0.828 ± 0.027 | 0.738 ± 0.078 |
| Fixed ($g = 0.5$) | 0.836 ± 0.036 | 0.822 ± 0.030 | 0.710 ± 0.082 |
| Open ($g = 1$) | 0.822 ± 0.035 | 0.816 ± 0.026 | 0.792 ± 0.079 |
| Random | 0.822 ± 0.039 | 0.802 ± 0.035 | 0.729 ± 0.084 |
| **Closed ($g = 0$)** | **0.553 ± 0.045** | **0.494 ± 0.047** | **1.124 ± 0.071** |

**Table 16.** Stream-intervention controls on SAAINT-DB using corrected nested-validation checkpoints. Each intervention is evaluated on the corresponding true outer held-out fold.

| Condition | Pearson $r$ | Spearman $\rho$ | RMSE |
| --- | --- | --- | --- |
| Full model | 0.838 ± 0.033 | 0.828 ± 0.027 | 0.738 ± 0.078 |
| Zero light | 0.764 ± 0.037 | 0.744 ± 0.036 | 0.889 ± 0.121 |
| Zero heavy | 0.592 ± 0.037 | 0.573 ± 0.022 | 1.048 ± 0.059 |
| Zero antigen | 0.593 ± 0.071 | 0.558 ± 0.067 | 1.057 ± 0.102 |
| Mean antigen | 0.510 ± 0.081 | 0.479 ± 0.083 | 1.199 ± 0.177 |
| Shuffled antigen | 0.365 ± 0.056 | 0.330 ± 0.076 | 1.330 ± 0.071 |

**Table 17.**
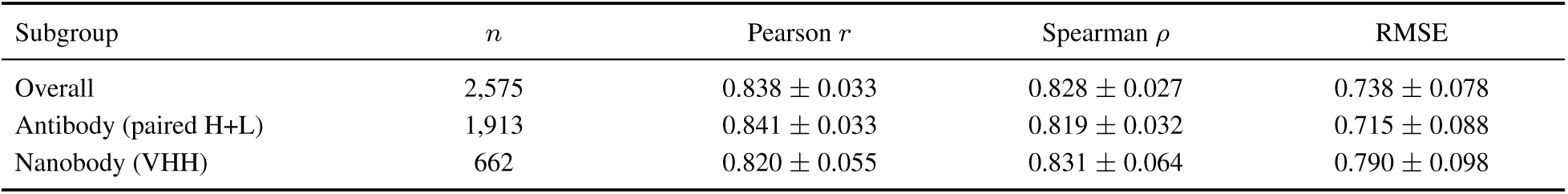
Performance stratified by paired antibodies and nanobodies on SAAINT-DB using corrected nested-validation checkpoints. Values are mean ± s.d. across the corrected evaluation folds, stratified by antibody format within each held-out fold.

**Table 18.**
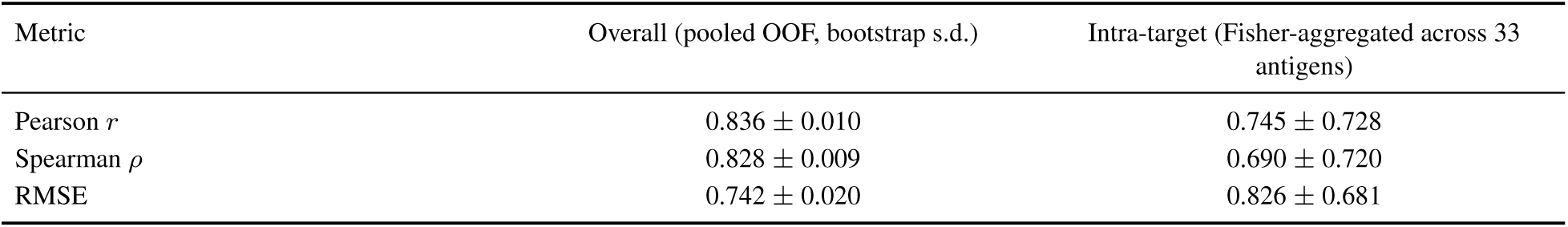
Overall versus intra-target ranking. Overall values are pooled out-of-fold metrics with bootstrap s.d.; intra-target values are Fisher-aggregated mean ± s.d. across 33 antigens.

| Metric | Overall (pooled OOF, bootstrap s.d.) | Intra-target (Fisher-aggregated across 33 antigens) |
| --- | --- | --- |
| Pearson $r$ | 0.836 ± 0.010 | 0.745 ± 0.728 |
| Spearman $\rho$ | 0.828 ± 0.009 | 0.690 ± 0.720 |
| RMSE | 0.742 ± 0.020 | 0.826 ± 0.681 |

## 13 Supplementary mutation and transfer analyses

These tables report heavy-chain CDR mutation performance, zero-shot to few-shot transfer and Top-K binder recovery for affinity-maturation use cases represented by AbBiBench and FLAb2-style landscapes (Zha et al., 2025; Chungyoun and Gray, 2025). Zero-shot transfer is informative but uneven across landscapes, whereas few-shot fine-tuning shows that the pretrained representation can be adapted with limited target-specific labels. Together, these analyses support use of AbAffinity as a triage system for prioritising variants and reducing experimental screening burden.

**Table 19.**
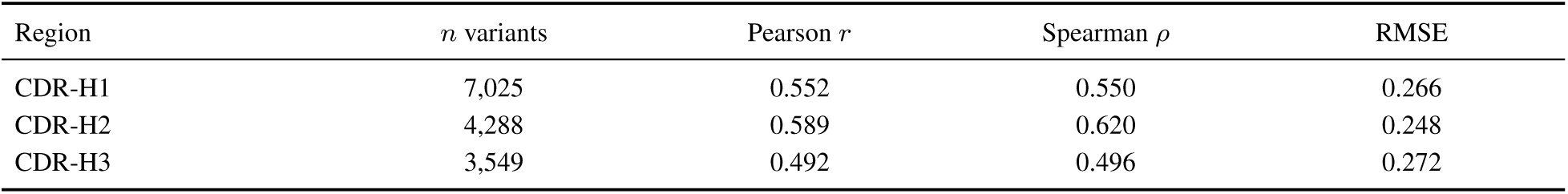
Per-CDR-region performance for heavy-chain single mutations. Single evaluation per region; no fold s.d.

| Region | *n* variants | Pearson *r* | Spearman $\rho$ | RMSE |
| --- | --- | --- | --- | --- |
| CDR-H1 | 7,025 | 0.552 | 0.550 | 0.266 |
| CDR-H2 | 4,288 | 0.589 | 0.620 | 0.248 |
| CDR-H3 | 3,549 | 0.492 | 0.496 | 0.272 |

**Table 20.** Transfer from zero-shot to few-shot fine-tuning with 30% labelled data. Few-shot uses validation-based early stopping with no test-set selection and affine range calibration.

| Dataset | Zero-shot Pearson | Zero-shot Spearman | @30% Pearson | @30% Spearman |
| --- | --- | --- | --- | --- |
| 1mlc | 0.137 | 0.166 | 0.428 ± 0.049 | 0.415 ± 0.036 |
| 1n8z | -0.316 | -0.351 | 0.690 ± 0.026 | 0.696 ± 0.044 |
| 4fqi | 0.452 | 0.469 | 0.967 ± 0.002 | 0.950 ± 0.008 |

**Table 21.** Top-K binder recovery on AAYL51. Values are fractions of experimentally top-ranked binders captured within each method’s own top-K% predictions.

| Cut-off | Random | Zero-shot | Mean-pool | AbAffinity |
| --- | --- | --- | --- | --- |
| Top-5% | 0.050 | 0.047 | 0.233 | **0.233** |
| Top-10% | 0.100 | 0.081 | 0.256 | **0.267** |
| Top-20% | 0.200 | 0.180 | 0.355 | **0.407** |
| Top-30% | 0.300 | 0.328 | 0.502 | **0.517** |

## 14 Supplementary structural attribution panels

The main text shows the complete heatmap-plus-structure comparison for 1VFB (Bhat et al., 1994), contrasting AbAffinity with the fused two-stream baseline. Supplementary attribution panels are provided below for 4ETQ, 5GRJ and 5Y9J (Matho et al., 2012; Liu et al., 2017; Shin et al., 2018). Integrated Gradients is evaluated as a spatial attribution method: high-scoring residues are compared with structurally validated interface positions, but the scores are not interpreted as residue-level ΔΔ*G* values or as a substitute for computational alanine scanning.

**Table 22.** Paratope recovery across normalised-attribution thresholds. Recovery is measured over 73 structurally validated heavy- and light-chain interface positions.

| Attribution threshold | AbAffinity | Two-stream baseline |
| --- | --- | --- |
| > 0.3 | 64/73 (88%) | 32/73 (44%) |
| > 0.4 (main text) | 56/73 (77%) | 22/73 (30%) |
| > 0.5 | 47/73 (64%) | 9/73 (12%) |
| > 0.6 | 26/73 (36%) | 4/73 (5%) |

**Figure 10.**
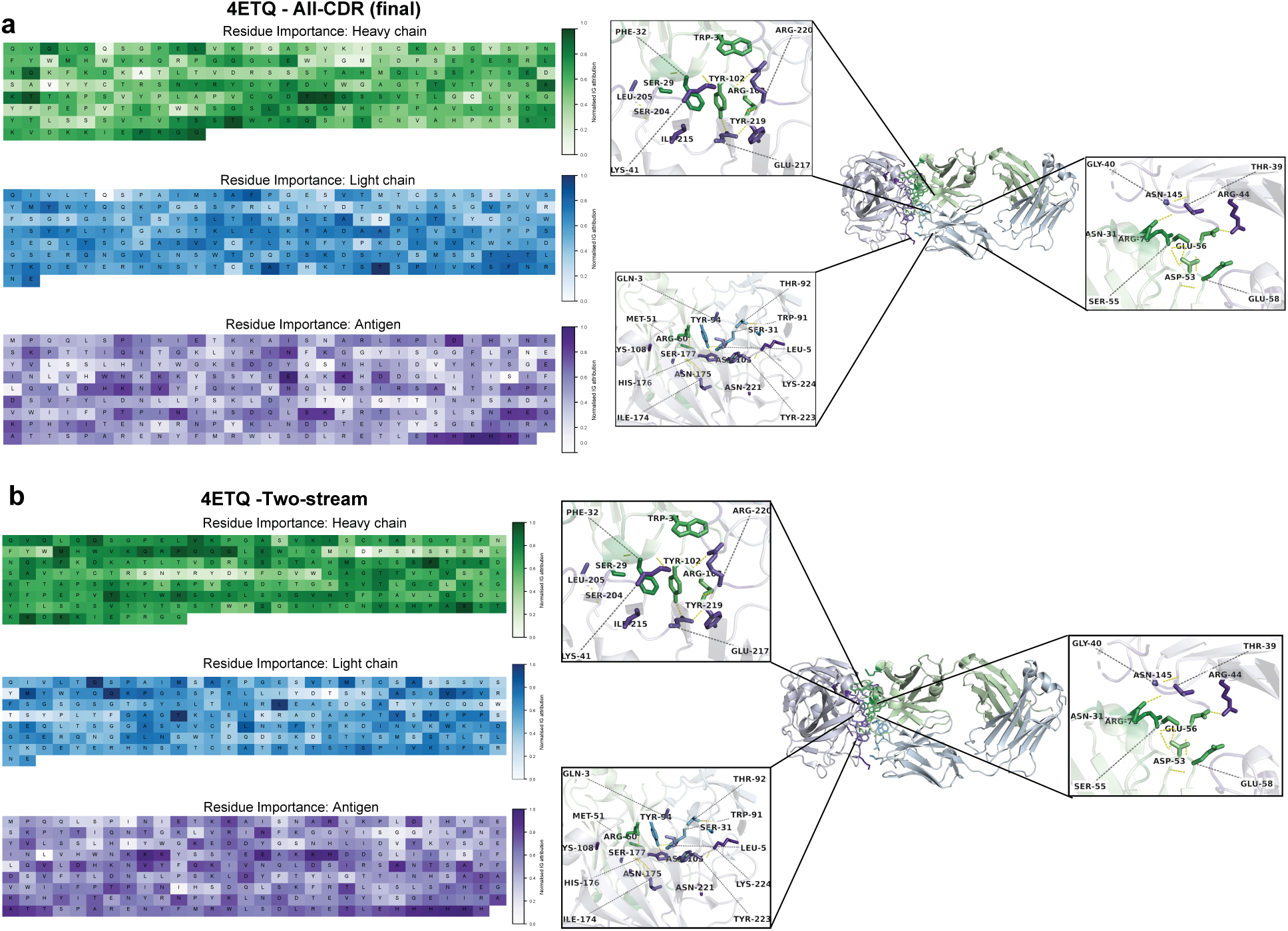
Structural attribution panel for PDB 4ETQ. Ground-truth interface residues were referenced from the LA5-D8 structural study (Matho et al., 2012). AbAffinity highlights CDR-proximal antibody residues and D8 epitope residues that coincide with the reported LA5-D8 contact network. In the structural zooms, high-attribution antibody residues include LA5 aromatic and polar CDR positions such as Phe32, Trp34, Ser29, Tyr94, Tyr102 and Arg103, while high-attribution D8 residues include Arg220, Tyr219, Glu217, Asn221, Lys224 and neighbouring surface residues. These positions are important in the ground-truth structure because the antibody uses aromatic CDR contacts and polar side chains to grip the D8 surface, with Arg/Glu/Asn-containing D8 patches contributing hydrogen-bond and electrostatic contacts. The two-stream baseline is more diffuse, whereas AbAffinity concentrates attribution on the CDR-D8 interface.

**Figure 11.**
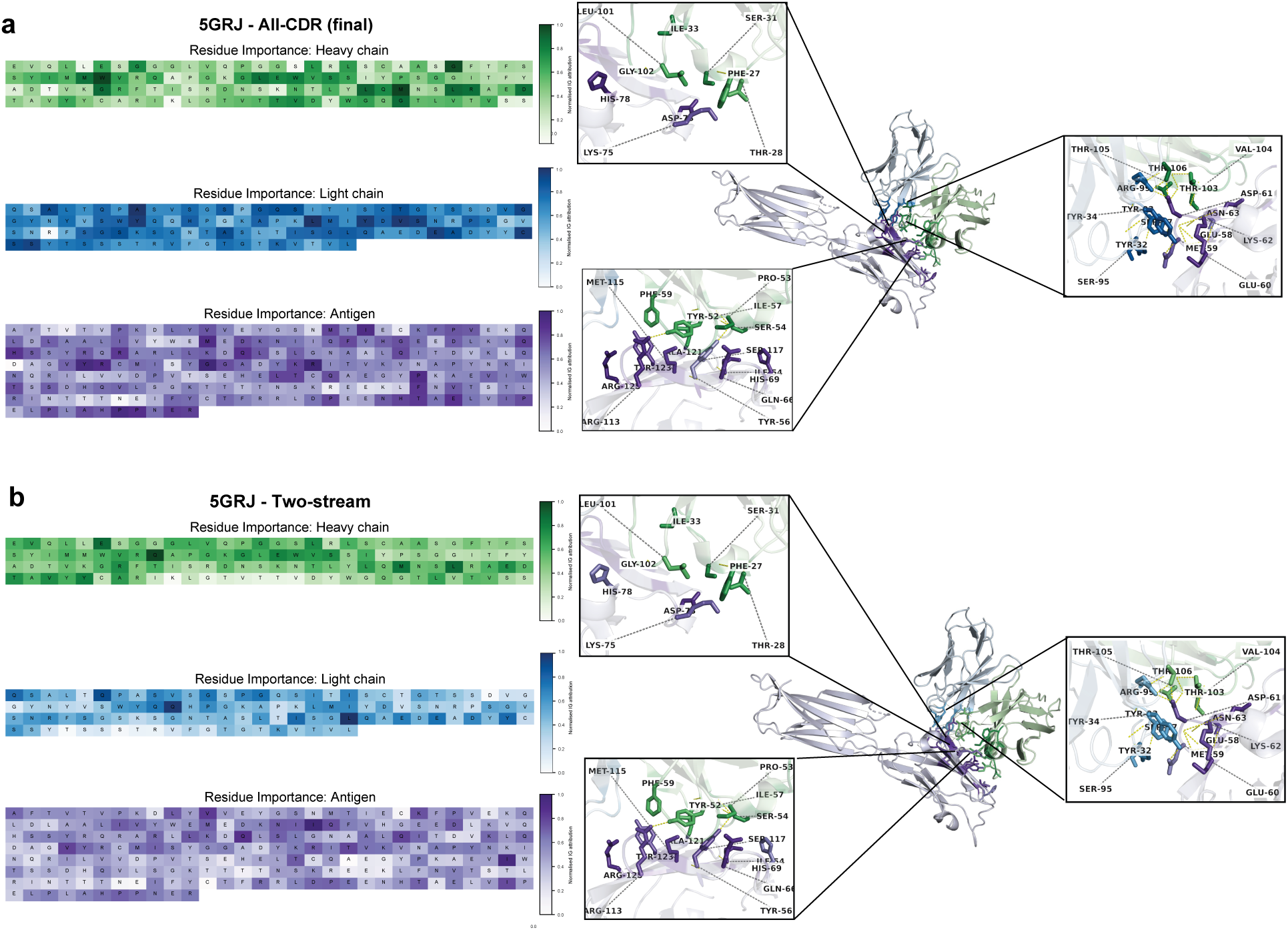
Structural attribution panel for PDB 5GRJ. Ground-truth interface residues were referenced from the avelumab-PD-L1 structural study (Liu et al., 2017). AbAffinity highlights residues on the avelumab CDR surface and the PD-L1 IgV-domain face used for checkpoint blockade. In the structural insets, attributed antibody residues include Tyr32/Tyr34, Arg99, Thr103/Val104/Thr105/Thr106 and nearby light-chain positions, while attributed PD-L1 residues include Glu58, Met59, Glu60, Asp61, Lys62, Asn63, Ser95, Met115, Ser117, Ala121, Arg123 and Tyr123-adjacent contacts. These residues are important because the avelumab epitope overlaps the PD-1 binding surface on PD-L1; acidic and polar PD-L1 residues around Glu58-Asn63 and the Met115-Arg123 region form the contact face that avelumab occupies to block receptor engagement. Compared with the two-stream baseline, AbAffinity places sharper signal on this receptor-blocking epitope and its opposing CDR residues.

**Figure 12.**
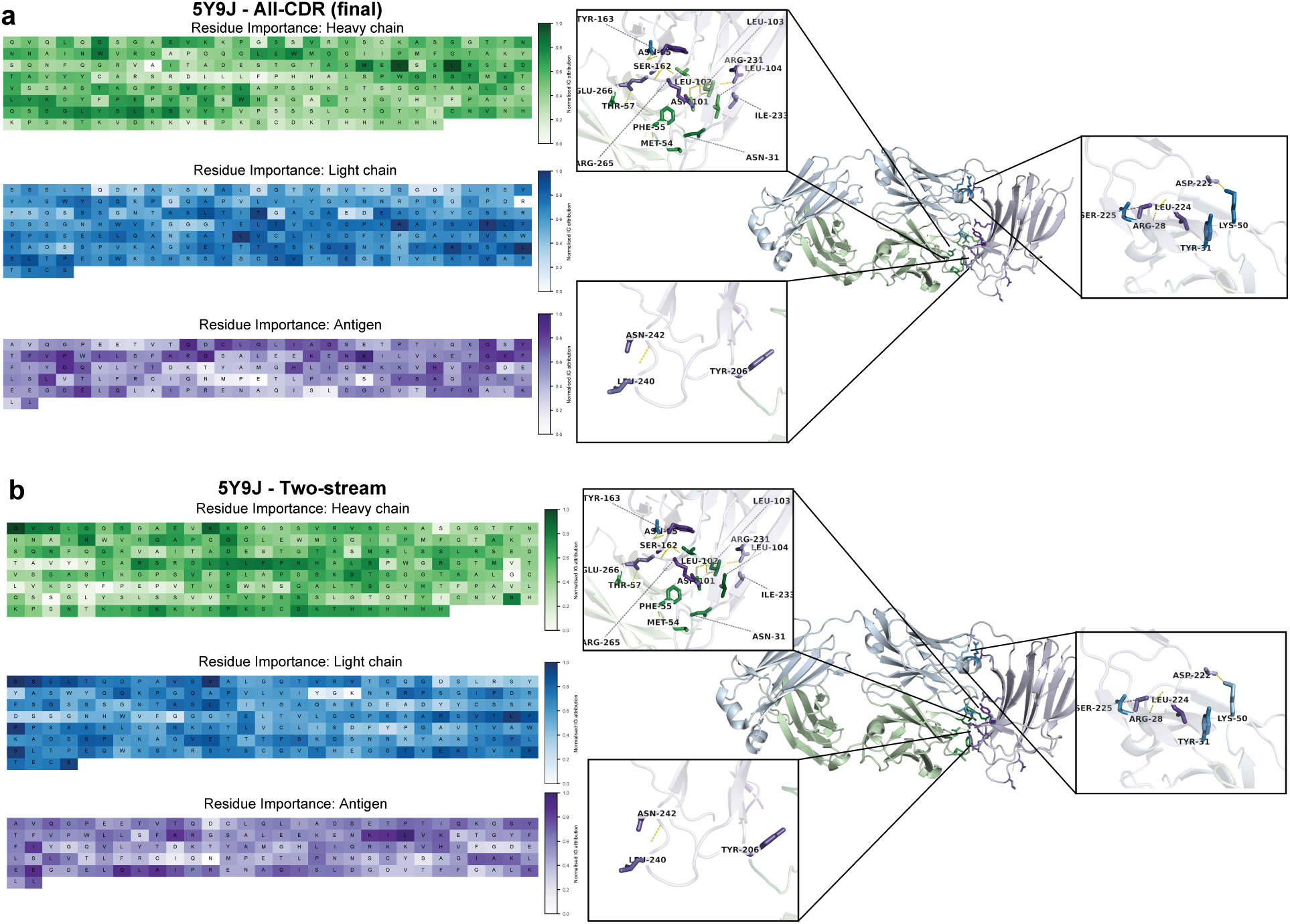
Structural attribution panel for PDB 5Y9J. Ground-truth interface residues were referenced from the belimumab-BAFF structural study (Shin et al., 2018). AbAffinity highlights belimumab CDR residues and BAFF residues on the receptor-binding surface reported in the structural study. The structural zooms show signal around BAFF Tyr163, Tyr206, Asp222, Leu224, Arg231, Ile233, Leu240, Asn242, Arg265 and Glu266, together with opposing antibody residues such as Met54, Phe55, Thr57, Asn101, Leu102/Leu103/Leu104, Ser162 and neighbouring CDR positions. These BAFF residues were selected in the ground-truth work because they define the belimumab-binding/receptor-overlap surface; aromatic and hydrophobic residues such as Tyr163, Tyr206, Leu224 and Leu240 help shape the binding patch, whereas charged or polar residues such as Asp222, Arg231, Asn242, Arg265 and Glu266 contribute specificity through hydrogen-bond and electrostatic interactions. AbAffinity therefore recovers residues that explain belimumab neutralisation of BAFF, while the two-stream baseline distributes attribution more broadly.

## Notes

https://github.com/harshitsinghsnu/AbAffinity

https://zenodo.org/records/23044968

### Competing Interest Statement

The authors have declared no competing interest.

### Summary of Updates

The Abstract and Introduction have been tightened to more clearly state the practical motivation for sequence-only affinity prediction and the specific limitations of existing sequence-based approaches. Methods Section 2 has been clarified. The description of heavy-chain CDR-focused pooling, heavy-light self-attention, the adaptive fusion gate, and the gated cross-attention blocks has been expanded with more precise equations and implementation details. The training and inference procedure for mapping cosine similarity to pKd is now stated more explicitly. Results have been reorganised for clarity. Nested random cross-validation, PDB-disjoint, and antigen-cluster-disjoint performance are presented together so that the contribution of related-antigen similarity is transparent. Matched baseline comparisons (concat+MLP and fused two-stream) and the CDR-pooling ablation are reported with consistent nested-validation checkpoints. Partner-shuffling, gate-intervention, and stream-ablation controls are expanded to support the claim of partner-conditioned prediction. Zero-shot and few-shot transfer results on affinity-maturation landscapes are presented with clearer interpretation of when target-specific calibration is required. The Discussion now more explicitly acknowledges the main generalisation gap (performance drop on antigen-sequence-divergent splits), the limited numerical benefit of the adaptive gate for conventional paired antibodies, and the retrospective nature of the current evidence. Prospective experimental validation is framed as the next required step. Figure 1 has been updated for visual clarity. Supplementary tables and figures have been expanded to include additional dataset statistics, per-format performance, and attribution recovery numbers. Author affiliations and corresponding-author contact details remain unchanged.

